# Population Genomics Reveals Molecular Determinants of Specialization to Tomato in the Polyphagous Fungal Pathogen *Botrytis cinerea* in France

**DOI:** 10.1101/2020.07.24.219691

**Authors:** Alex Mercier, Adeline Simon, Nicolas Lapalu, Tatiana Giraud, Marc Bardin, Anne-Sophie Walker, Muriel Viaud, Pierre Gladieux

## Abstract

Many fungal plant pathogens encompass multiple populations specialized on different plant species. Understanding the factors underlying pathogen adaptation to their hosts is a major challenge of evolutionary microbiology, and it should help preventing the emergence of new specialized pathogens on novel hosts. Previous studies have shown that French populations of the grey mould pathogen *Botrytis cinerea* parasitizing tomato and grapevine are differentiated from each other, and have higher aggressiveness on their host-of-origin than on other hosts, indicating some degree of host specialization in this polyphagous pathogen. Here, we aimed at identifying the genomic features underlying the specialization of *B. cinerea* populations to tomato and grapevine. Based on whole genome sequences of 32 isolates, we confirmed the subdivision of *B. cinerea* pathogens into two genetic clusters on grapevine and another, single cluster on tomato. Levels of genetic variation in the different clusters were similar, suggesting that the tomato-specific cluster has not recently emerged following a bottleneck. Using genome scans for selective sweeps and divergent selection, tests of positive selection based on polymorphism and divergence at synonymous and non-synonymous sites and analyses of presence/absence variation, we identified several candidate genes that represent possible determinants of host specialization in the tomato-associated population. This work deepens our understanding of the genomic changes underlying the specialization of fungal pathogen populations.

## Introduction

Many fungal plant pathogens encompass multiple lineages, host races or *formae speciales* specialized on different plant species. Understanding the proximate (*i.e.* molecular) and ultimate (*i.e.* eco-evolutionary) factors underlying adaptation to hosts is a major goal for evolutionary microbiology, because emerging diseases are often caused by the appearance and spread of new pathogen populations specialized onto new hosts (Fisher *et al.* 2012; Stukenbrock & McDonald 2008). Evolutionary theory predicts that pathogen specialization should facilitate the emergence of new populations onto novel hosts, because specialization restricts encounters of potential mates within hosts and reduces the survival of offspring due to maladaptation of immigrants and hybrid offspring, thereby reducing gene flow between ancestral and emerging populations (Giraud *et al.* 2010; Nosil *et al.* 2005). The role of specialization as a barrier to gene flow is expected to be strong for pathogens mating within or onto their hosts because, for individuals evolving the ability to infect novel hosts, mating automatically becomes assortative with respect to host use, and reproductive isolation arises as a direct consequence of adaptive divergence (Gladieux *et al.* 2011; Servedio *et al.* 2011). Evolutionary theory also predicts that specialization, and the associated emergence of new populations, could be facilitated by the molecular basis of plant-pathogen interactions, because compatibility is often determined by a limited number of genes in the host and the pathogen, and selection is more efficient when it acts on a smaller number of genes (Giraud *et al.* 2010; Schulze-Lefert & Panstruga 2011). However, despite the apparent ubiquity of specialized fungal pathogens and the negative impact that specialized populations can have on food security and ecosystem health, the genomic features involved in host specialization remain largely unknown. Acquiring knowledge about the genomic features underlying pathogen specialization can provide key insights into the mechanisms of specialization.

The ascomycete *Botrytis cinerea* is often presented as a textbook example of a polyphagous plant pathogen, parasitizing more than 1400 host plant species belonging to 580 genera (Elad *et al.* 2016). Previous population genetic studies reveal that *B. cinerea* is not strictly speaking a generalist pathogen, and that a more appropriate qualifier is “polyspecialist”, i.e. a set of populations specialized to different host species. Indeed, with the exception of California (Ma and Michailides, 2005; Caseys et al, 2020; Soltis et al, 2019; Atwell et al, 2015), studies conducted in multiple regions of the world reveal that populations were structured (reviewed in Walker 2016), and recognize the host as the factor with the highest explanatory power for population structure in *B. cinerea*, ahead of geography. In France, our previous work revealed population subdivision in *B. cinerea*, with genetic differentiation between field populations infecting tomato (*Solanum lycopersicum*) and grapevine (*Vitis vinifera*), respectively (Walker *et al.* 2015). This population structure in *B. cinerea* was shown to be stable in time and was observed in multiple regions in France. Furthermore, this structure was later associated with differences in performance on the two hosts, with pathogens isolated from tomato being more aggressive on tomato than pathogens isolated from grapevine, and reciprocally (Mercier *et al.* 2019). Altogether, these data were consistent with a certain degree of specialization of *B. cinerea* populations onto these two host plants.

Here, we aimed to identify the molecular basis of host specialization in the *B. cinerea* tomato- and grapevine-associated populations, by addressing the following questions: (1) Can we confirm the genetic subdivision between *B. cinerea* populations from tomato and grapevine using genomic data, and what is the degree of divergence between them? (2) Can we identify genes with footprints of positive selection and/or divergent selection in the genomes of populations specialized to different hosts, and what are their predicted functions? (3) Is there variation in gene content between *B. cinerea* populations associated with tomato and grapevine? To address these questions, we used a set of *B. cinerea* isolates collected on tomato and grapevine in different regions of France. We Illumina-sequenced their genomes and identified single nucleotide polymorphisms by mapping sequencing reads against a high-quality reference genome (van Kan *et al.* 2017). Because some tests of selection can be biased by population subdivision, while other tests are based on patterns of population differentiation, we first analyzed the population structure of *B. cinerea* collected on tomato and grapevine. To detect genes potentially involved in the specialization of *B. cinerea* to tomato, we searched for signatures of positive selection, by scanning genomes in the tomato population for selective sweeps, and by estimating the direction and intensity of selection using McDonald-Kreitman tests on coding sequences. Furthermore, we investigated signatures of divergent selection using genomic differentiation between populations, and we characterized variations in the presence/absence of predicted genes between populations collected on tomato and grapevine using *de novo* genome assemblies, gene prediction, and orthology analysis.

## Materials and methods

### Sample collection

*Botrytis cinerea* samples were selected in a collection of isolates (i.e. single-spored mycelial colonies) originating from three regions of France (Champagne, Occitanie and Provence) and previously characterized using analyses of population structure based on microsatellite markers and pathogenicity tests (Walker *et al.* 2015; Mercier et al., 2019). For each region and each host, we randomly selected three to nine isolates with high membership proportions (*q*>0.9) in the cluster matching their host of origin in a previous analysis of population structure based on microsatellite genotyping (Table 1; Mercier *et al.* 2019). Collection sites were 15 to 133 km apart within regions, and 204 to 722 km apart between regions (Supplementary Figure S1). The set of 32 isolates originated from the following hosts: (i) tomato (*Solanum lycopersicum*; fruits; 13 isolates), (ii) grapevine (*Vitis vinifera*; berries; 16 isolates), (iii) bramble (*Rubus fruticosus*; berries; two isolates) and (iv) hydrangea (*Hydrangea macrophylla*; flower buds; one isolate). Samples from tomato originated from plastic tunnels with sides opened (Occitanie region) or glasshouses (Provence and Champagne regions). Mycelia were cultured on malt-yeast-agar (MYA; 20 g.L^−1^ malt extract, 5 g.L^−1^ yeast extract, 15 g.L^−1^ agar) at 23°C under continuous light until conidiation, and stored as conidial suspensions in glycerol 20% at −80°C until use.

**Table 1.**
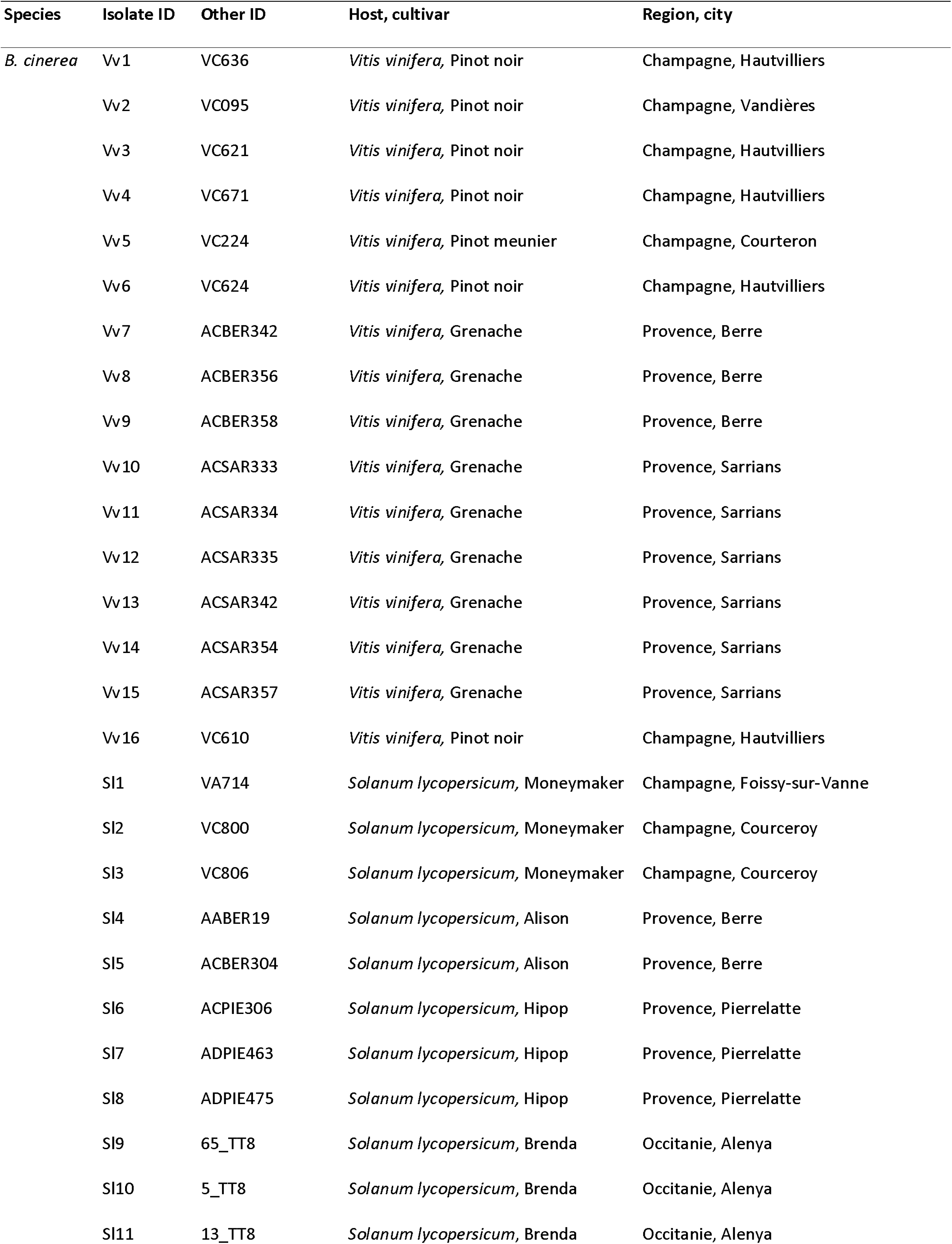
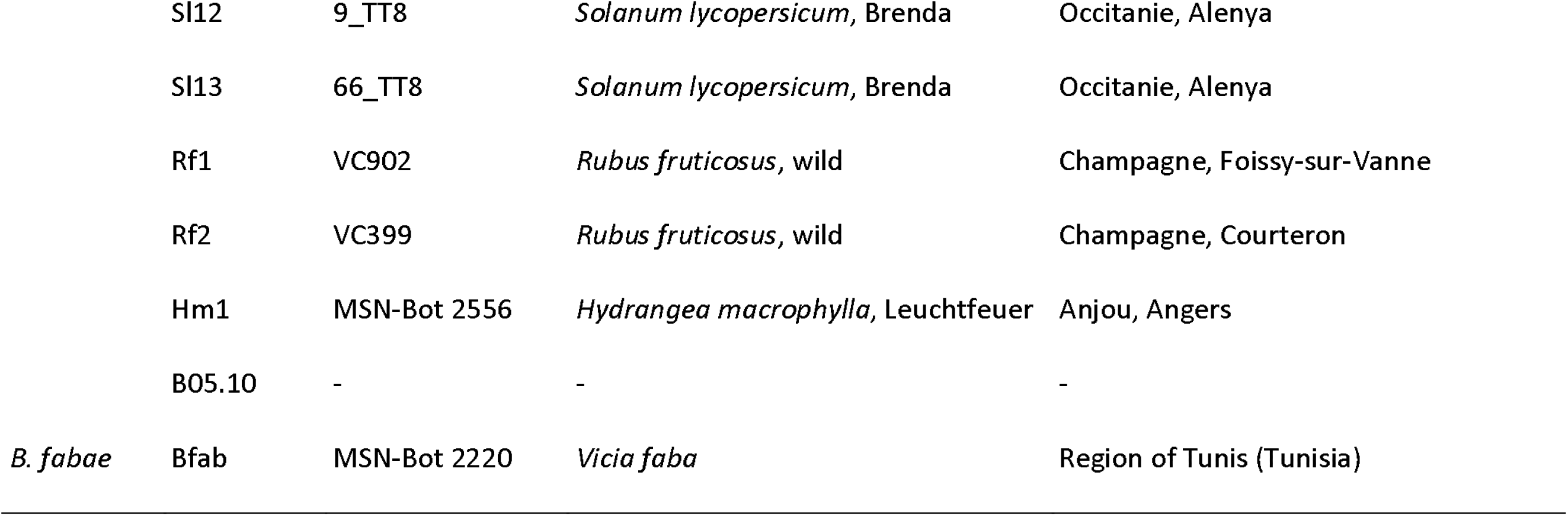
*Botrytis* isolates included in the study. Isolates from grapevine berries and tomatoes were collected in three regions of France between September 2005 and June 2007 for Champagne and Provence isolates (Walker *et al.* (2015), and between May and June 2009 for Occitanie isolates. Tomatoes in Occitanie were grown in plastic tunnels with sides open, whereas those from the Champagne and Provence areas were grown in glass-greenhouses. B05.10 is used as the reference isolate for genomic analysis (van Kan *et al.* 2016)

### Pathogenicity tests on tomato plants

Isolates of *B. cinerea* collected from tomato or grape were cultivated on MYA medium in a growth chamber (21°C, 14 hours light) for 14 days. Conidia were then washed with sterile distilled water. The conidial suspension was filtered through a 30 µm mesh sterile filter to remove mycelium fragments. The conidial concentration was determined with a haemacytometer and adjusted to 10^6^ spores/mL. Seeds of tomato var. Clodano (Syngenta) were sown in compost and transplanted after one week in individual pots. Plants were grown in a glasshouse for 6 to 8 weeks where they received a standard commercial nutrient solution once or twice a day, depending on needs. Plants had at least eight fully expanded leaves when inoculated with a conidial suspension. Each isolate was inoculated on five plants, from each of which two leaves were removed, leaving 5-10 mm petiole stubs on the stems. Each pruning wound was inoculated with 10 µl of conidial suspension at 10^6^ conidia/mL. Inoculated plants were incubated in a growth chamber in conditions conducive to disease development (21°C, 16h-photoperiod, 162 µmol.s^−1^ .m^−2^, relative humidity > 80%). Due to a growth chamber area that did not allow all the isolates to be tested at the same time, nine series of pathogenicity tests were conducted with 10-12 isolates each, together with the BC1 reference isolate collected in 1989 in a tomato glasshouse in Brittany (Decognet *et al.* 2009). For each isolate, two to four independent repetitions of the pathogenicity test were performed. Lesion sizes (in mm) were assessed daily between the 4^th^ and the 7^th^ day post-infection and the Area Under the Disease Progress Curve (AUDPC; (Simko & Piepho 2012) was computed to take into account the kinetics of disease development for each isolate. To compare the aggressiveness of isolates, an aggressiveness index (AI), relative to the reference isolate BC1, was computed as follows: *AI_isolate_* = 100 × (*AUDPC_isolate_/AUDPC*_*BC*1_), where *AUDPC_isolate_* was the average AUDPC for a given isolate and *AUDPC_BC1_* is the average AUDPC for the reference isolate BC1. Using the AI index calibrating the AUDPC of a given isolate with that of the reference isolate BC1 in the same test allows comparing isolates aggressiveness while taking into account the variability occurring among assays (e.g. plant physiological state; (Leyronas *et al.* 2018). Because of data non-normality, data were analysed using non-parametric tests. Three statistical tests were carried out with Statistica: (1) we used a non-parametric analysis of variance (Kruskal-Wallis test) to assess differences among isolates in terms of aggressiveness on grape and tomato, considering the average values for each of the independent pathogenicity tests as replications; (2) we used the Mann-Whitney U-test to compare the aggressiveness of isolates from different hosts of origin (tomato vs. grape), considering the independent repetitions of the pathogenicity test as blocks and the 13 isolates from tomato and 16 isolates from grapevine as replicates; (3) we used a Kruskal-Wallis test to compare isolates from three different clusters (see *Results* section) in terms of aggressiveness on tomato plants.

### DNA preparation and sequencing

Isolates were cultivated for 48 h on MYA + cellophane medium at 23 °C in the dark and then ground using a mortar and pestle in liquid nitrogen. DNA was extracted using a standard sarkosyl procedure (Dellaporta *et al.* 1983). Paired-end libraries were prepared and sequenced (2 x 100 nucleotides) on a HiSeq4000 Illumina platform at Integragen (Evry, France). Sequencing coverage ranged from 58 to 305 X. Genomic data were deposited at SRA under accession number PRJNA624742. Read quality was checked using FastQC (https://www.bioinformatics.babraham.ac.uk/projects/fastqc/).

### SNP calling and filtering

SNPs were detected with the same workflow as described in (Zhong *et al.* 2017). Sequencing reads were preprocessed with Trimmomatic v0.36 (Bolger *et al.* 2014). Preprocessed reads were mapped onto the *B. cinerea* B05.10 reference genome (van Kan *et al.* 2017) using BWA v0.7.15 (Li & Durbin 2009). Aligned reads were filtered based on quality using SamTools v1.3 (Li *et al.* 2009) and Picard Tools (http://broadinstitute.github.io/picard/) to remove secondary alignments, reads with a mapping quality <30 and paired reads not at the expected distance. SNP calling was performed with Freebayes v1.1 (Garrison & Marth 2012). Further filtering was carried out using script VCFFiltering.py (https://urgi.versailles.inra.fr/download/gandalf/VCFtools-1.2.tar.gz), following (Li 2014). We kept only biallelic SNPs supported by more than 90% of aligned reads, detected outside low-complexity regions or transposable elements (as identified in the reference isolate B05.10: https://doi.org/10.15454/TFYH9N; Porquier *et al.* 2016) and with coverage lower than twice the standard deviation from the mean depth coverage. The VCF file is available on Zenodo (doi: 10.5281/zenodo.4293375).

### Population structure and demographic history

We performed a principal component analysis based on biallelic SNPs using the python library scikit-allel 1.3.2 (https://github.com/cggh/scikit-allel). We used the SNMF program to infer individual ancestry coefficients in K ancestral populations. This program is optimized for the analysis of large datasets and it estimates individual admixture coefficients based on sparse non-negative matrix factorization, without assuming Hardy-Weinberg equilibrium (Frichot et al., 2014). We used Splitstree 4 (Huson & Bryant 2005) to visualize relationships between genotypes in a phylogenetic network, with reticulations representing the conflicting phylogenetic signals caused by recombination or incomplete lineage sorting. The position of the root was determined using a *B. fabae* isolate as the outgroup. Botrytis fabae is one of the closest known relatives of *B. cinerea* (Amselem *et al.* 2011; Walker 2016). Summary statistics of genomic variation (segregating sites S, nucleotide diversity π, Watterson’s θ, Tajima’s D) were estimated using EGGLIB 3.0 (https://egglib.org/) on 10kb windows, excluding sites with more than 50% missing data and removing windows with lseff<1000 (lseff is the number of analysed sites) or nseff<3 (nseff is the average number of exploitable samples). Site frequency spectra were estimated using dadi 1.7.0 (Gutenkunst et al. 2009). Allelic richness and private allele richness were estimated with ADZE 1.0 (Szpiech *et al.* 2008), using a generalized rarefaction approach to account for differences in sample size among populations.

We employed the *f3* statistic (Reich *et al.* 2009) to test for admixture based on shared genetic drift, as implemented in the Popstats python script (Skoglund *et al.* 2015; https://github.com/pontussk/popstats/). The *f3* statistic is used to test for admixture among three populations Px, P1, P2. In the no-admixture case, the *f3* statistic measures the branch length between Px and the internal node of the unrooted population tree (Px;P1,P2), and it is therefore expected to be greater than zero. In the case Px has a mixed ancestry from P1 and P2, or populations closely related to them, the *f3* statistic is expected to be negative. Significance was assessed by block jackknife by treating each chromosome as a block and weighting each block by the number of SNPs. The standard error of the test statistic was used to define a Z-score.

### Genome assembly and gene prediction

Gene content was determined using two independent approaches. In the first approach, we used the read mapping coverage of the genes previously predicted in the reference genome B05.10 (van Kan *et al.* 2017). Read count per site for each gene and for each isolate was computed using SAMTOOLS BEDCOV, and normalized using edgeR (McCarthy *et al.* 2012; Robinson *et al.* 2010). Genes showing significant differences in mapping coverage across populations were identified with edgeR (adjusted p-value <0.05 and more than two-fold change). Putative gene duplication events (higher mapping coverage) and missing B05.10 genes (lack of mapping reads) were visually inspected in their genomic context using a genome browser for validation.

The second approach was based on genome assembly and *de novo* gene prediction. Illumina paired-reads were assembled using a combination of Velvet (Zerbino & Birney 2008), SOAPDENOVO and SOAPGAPCLOSER (Luo *et al.* 2012), as follows: (1) reads were trimmed at the first N, (2) contigs were generated with several k-mer values using SOAPDENOVO, (3) several Velvet assemblies were built using several k-mer values and as the input the trimmed reads and all SOAPDENOVO contigs considered as “long reads”, (4) the assembly that maximizes the criterion (N50*size of the assembly) was selected, (5) SOAPGAPCLOSER was run on the selected assembly, and (6) Contigs completely included in other longer contigs were deleted. Genomic regions mapping to transposable elements previously identified in *B. cinerea* (https://doi.org/10.15454/TFYH9N; Porquier *et al.* 2016) were masked with RepeatMasker (http://www.repeatmasker.org) prior to gene prediction. Genes were predicted using the FGenesh *ab initio* gene-finder (Solovyev *et al.* 2006; http://www.softberry.com/berry.phtml), the program previously used to annotate the reference genome (Amselem *et al.* 2011), and for which *Botrytis*-specific gene-finding parameters were thus available. Completeness of the assembly and gene prediction were evaluated with BUSCO using the Ascomycota gene set (Seppey *et al.* 2019). We then used OrthoFinder (Emms & Kelly 2015) in order to define groups of orthologous sequences (hereafter “groups of orthologs”) based on sequences of predicted genes translated into protein sequences. To reduce the impact of incomplete gene prediction (e.g. truncated genes in small contigs), groups of orthologs were then manually checked for the presence/absence of the protein-encoding genes in the genomes using TBLASTN (Johnson *et al.* 2008). Proteins were functionally annotated using InterProScan (Jones *et al.* 2014), and SignalP (Almagro Armenteros *et al.* 2019) was used to predict secretion signal peptides. Prediction of transmembrane helices in proteins was performed using TMHMM 2.0 (Krogh *et al.* 2001; Sonnhammer et al.). As we detected traces of bacterial contamination in the genome of the Sl13 isolate, this genome was not included in orthology analysis. However, we have kept this genome for analyzes based on polymorphism, because bacterial reads cannot map to the reference genome.

### Tests of positive selection based on polymorphism and divergence at synonymous and non-synonymous sites

We estimated the intensity and direction of selective pressures exerted on genes in populations using the McDonald-Kreitman test based on polymorphism and divergence at synonymous and non-synonymous sites (McDonald & Kreitman 1991). This test is based on the number of nucleotide polymorphisms and substitutions in gene sequences, and assumes that synonymous mutations are neutral. Pseudo-sequences for coding sequences of all genes in the reference genome were generated using the VCF file and the reference sequence. Synonymous and non-synonymous divergence was computed with CODEML (model 0; Yang 1997, 2007), using *B. fabae* as the outgroup. Synonymous and non-synonymous polymorphism was computed using EGGLIB 3 (https://egglib.org/), filtering out sites with more than 80% missing data and excluding filtered alignments with less than 10 codons or four sequences. Samples Sl1, Sl2, Sl3, Sl13, Vv3, Vv5, Vv2, Vv4 and Vv6 were excluded because they introduced missing data that prevented calculations of synonymous and non-synonymous divergence. The neutrality index (NI), defined as (P_N_/D_N_)/(P_S_/D_S_) (Stoletzki & Eyre-Walker 2011), was computed for every gene, with P_N_ and D_N_ the numbers of non-synonymous polymorphisms and substitutions, respectively, and P_S_ and D_S_ the numbers of synonymous polymorphisms and substitutions, respectively. Pseudocounts of one were added to each cell of the McDonald-Kreitman tables to ensure the NI was always defined (i.e. no division by zero).

### Linkage disequilibrium and recombination

We used PopLDdecay (Zhang *et al.* 2019a) to measure linkage disequilibrium (*r*^2^) as a function of the distance between pairs of SNPs. PopLDdecay was configured with a maximum distance between SNPs of 300 kbp, a minor allele frequency of 0.005 and a maximum ratio of heterozygous allele of 0.88. Recombination rates were estimated for each chromosome with PAIRWISE in LDHAT version 2.2 (Auton & McVean 2007). Sites with missing data were excluded.

### Tests of positive selection based on linkage disequilibrium and the site frequency spectrum

We searched for signatures of selective sweeps along genomes using three different softwares, each implementing a different approach. The SweeD 3.0 software (Pavlidis *et al.* 2013) implements a composite likelihood ratio (CLR) test based on the SweepFinder algorithm (Nielsen *et al.* 2005), which uses the site frequency spectrum (SFS) of a locus to compute the ratio of the likelihood of a hard selective sweep at a given position to the likelihood of a null model hypothesis without selection. The CLR statistic was computed for each chromosomes of each population using a grid size of 50 or 200 (grid size is the number of positions where the likelihood is calculated), but only results for a grid size of 200 are presented because the CLR profiles were highly similar between the two settings. Input files included both “unfolded” SNPs (i.e. SNPs for which ancestral and derived states can be determined using the allelic state of the outgroup) and “folded” SNPs (i.e. SNPs for which the outgroup had missing data). Only biallelic sites with sample size greater than or equal to five were included. The *nS_L_* method implemented in the nSL software (Ferrer-Admetlla *et al.* 2014) detects hard and soft selective sweeps based on haplotype homozygosity. The nSL statistic was computed for each chromosome of each population, including only biallelic sites with sample size greater than or equal to five. Nine isolates (Sl1, Sl2, Sl3, Sl13, Vv3, Vv5, Vv2, Vv4, Vv6) were excluded to reduce the proportion of missing data. The *hapFLK* method (Fariello *et al.* 2013) implemented in the HapFLK software is based on the original *FLK* method by (Bonhomme *et al.* 2010), which detects signatures of selection from differentiation between populations. This metric was used to test the null hypothesis of neutrality by contrasting allele frequencies at a given locus in different populations. *hapFLK* extends the FLK method to account for the haplotype structure in the sample, and the method is robust to the effects of bottlenecks and migration. HapFLK takes the number of cluster of haplotypes as a parameter (K). To determine the number of clusters of haplotypes K, we ran FASTPHASE v1.4 (Scheet & Stephens 2006) and R package IMPUTEQ (Khvorykh & Khrunin 2020) on polymorphism data for the largest chromosome BCIN01. For each population, we used IMPUTEQ to generate five datasets with 10% of polymorphic positions masked, and for each masked dataset we used FASTPHASE for imputing masked positions assuming clusters of K=2,3…10 haplotypes and the following parameters: -T10 -C25 -H-1 -n -Z. We estimated error using Estimate-Errors function in IMPUTEQ and the number of clusters that minimizes the error was selected as the optimum. The HapFLK metric was computed individually on each chromosome using as number of clusters K = 5, and nfit = 2. Accessory chromosomes were not included in these analyses as they showed presence/absence polymorphism (see *Results* section).

## Results

### Whole-genome sequencing, population structure and demographic history

Previous work using microsatellite data revealed differentiation between populations of *B. cinerea* collected on tomato and grapevine (Mercier *et al.* 2019; Walker *et al.* 2015). To investigate the genetic basis of specialization of the tomato- and grapevine-infecting populations, we randomly selected for genome sequencing 32 isolates collected on tomato (13 isolates), grapevine (16 isolates), *Rubus* (two isolates) and *Hydrangea* (one isolate) (Table 1). Isolates from *Rubus* and *Hydrangea* were previously shown to belong to generalist populations (i.e. assigned to clusters found on all sampled hosts; Mercier *et al.* 2019). One isolate of the sister species *B. fabae* was also sequenced and used as the outgroup. Information about the sequenced isolates is summarized in Table 1. Alignment of sequencing reads to the B05.10 reference genome (van Kan *et al.* 2017) followed by SNP calling identified 249,084 high-quality SNPs.

In clustering analyses based on sparse nonnegative matrix factorization algorithms, as implemented in the SNMF program (Frichot et al., 2014), the model with *K*=4 clusters was identified as the best model based on cross-entropy (Supplementary Figure S2) and models with K>4 did not identify well-delimited and biologically relevant clusters (Fig. 1A). At K=4, one cluster was associated with tomato (hereafter referred to as “T” cluster), two clusters were associated with grapevine (hereafter referred to as “G1” for the largest, and “G2” for the smallest), and one cluster was formed by the isolates Rf1 and Hm1 from bramble (*R. fruticosus*) and hydrangea (*H. macrophylla*) (Figure 1A). The isolate Rf2 collected on wild blackberry displayed ancestry in multiple clusters, and the reference isolate B05.10 had ancestry in the G2 and T clusters. No pattern of geographical subdivision was observed, consistent with previous findings (Walker *et al.* 2015; Mercier et al., 2019). The neighbor-net network inferred with Splitstree revealed three main groups, corresponding to the three main clusters identified with SNMF, with cluster G2 (isolates Vv8, Vv9, Vv11 and Vv15) more closely related to T than to G1 (Figure 1B). Reticulations in the neighbor-net network and patterns of membership at K=2 and K=3 in the analysis with sNMF indicated that cluster G2 shared recent ancestry with clusters G1 and T. However, tests for admixture using the *f3* statistic (Reich *et al.* 2009; Skoglund *et al.* 2015) did not support a scenario in which G2 derived from admixture between the two other clusters (Supplementary Table S1). The principal component analysis corroborated the results of clustering analyses and the neighbor-net network (Figure 1C). The first principal component separated isolates Hm1 and Rf1 from the rest of the dataset. The second principal component individualized isolate Rf2, as well as the three clusters T, G1 and G2. The third and fourth principal components individualized cluster G2 and isolate Sl11, respectively. Together, analyses of population subdivision revealed three clearly defined populations (two on grapevine and one on tomato) and we therefore focused on these populations to identify the genes underlying differences in host specialization.

**Figure 1.**
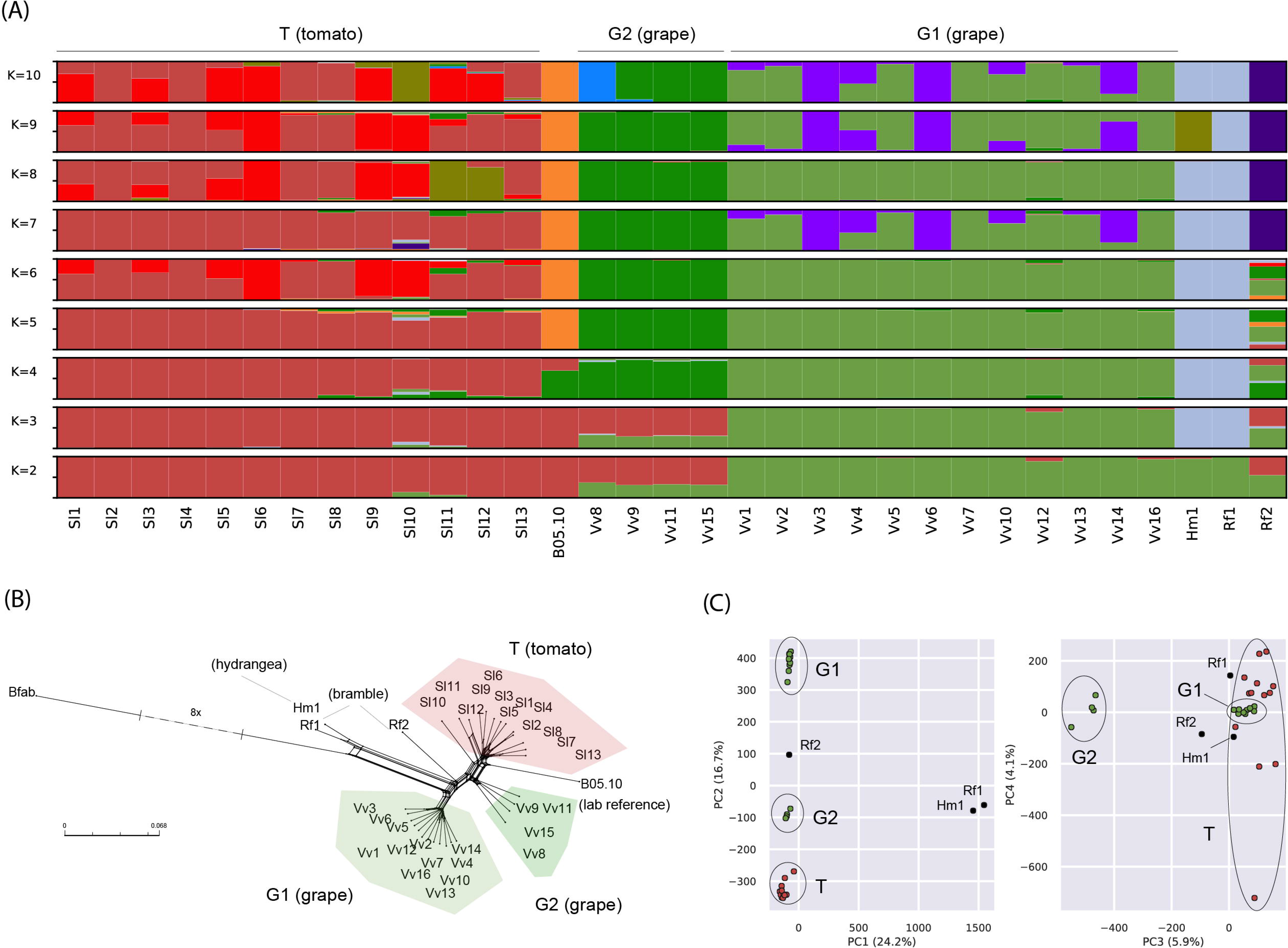
Population subdivision inferred based on SNPs identified in 32 isolates of *Botrytis cinerea* collected on tomato (red, Sl prefix, *Solanum lycopersicum*), grape (green, Vv prefix, *Vitis vinifera*), bramble (black, Rf prefix, *Rubus fruticosus*) and hydrangea (black, Hm prefix, *Hydrangea macrophylla*). (A) Ancestry proportions in K clusters, as estimated with the SNMF program. Each multilocus genotype is represented by a vertical bar divided into K segments, indicating membership in K clusters. (B) Neighbor-net phylogenetic network estimated with Splitstree, with one isolate of *B. fabae* (Bfab) used as the outgroup. Reticulations indicate phylogenetic conflicts caused by recombination or incomplete lineage sorting. (C) Principal component analysis showing first four principal components PC1, PC2, PC3 and PC4. Isolate B05.10 in (A) and (B) is the reference genome for *B. cinerea* (van Kan *et al.* 2016)

On average across core chromosomes BCIN01 to BCIN16, nucleotide diversity π and Watterson’s θ were comparable in the three populations (from π=0.0018/bp in G1 to π =0.0030/bp in G2; from θ=0.0017/bp in G1 to θ=0.0027/bp in G2; Table 2; Supplementary Table S2), although all comparisons were statistically significant except between populations T and G2 for θ (two-tailed Wilcoxon signed-rank test, P-value>0.05). Allelic richness was slightly higher in G2 than in T (AR=1.104 vs AR=1.094), and lower in G1 (AR=1.078). Private allele richness was higher in G1 (PAR=0.085), than in T (PAR=0.073) and G2 (PAR=0.071) (Table 2).

**Table 2.** Summary statistics of genomic variation in three clusters of *Botrytis cinerea*

| Cluster | S | $\pi$ | $\theta$ | AR | PAR | D | H | LD50 | $\rho$ |
| --- | --- | --- | --- | --- | --- | --- | --- | --- | --- |
| T | 37763 | 0.0027 | 0.0026 | 1.094<br>(0.0002) | 0.073<br>(0.0002) | 0.309 | 0.451 | 8600 | 0.0108 |
| G1 | 26769 | 0.0018 | 0.0017 | 1.078<br>(0.0002) | 0.085<br>(0.0002) | 0.167 | 0.230 | 3100 | 0.0178 |
| G2 | 14307 | 0.0030 | 0.0027 | 1.104<br>(0.0003) | 0.071<br>(0.0002) | 1.470 | 0.140 | 16800 | 0.2047 |

S, number of segregating sites; $\pi$ , nucleotide diversity per site; $\theta$ , Watterson's estimate of the population mutation parameter per site; $\rho$ , population recombination parameter per site; AR, allelic richness (standard error of the mean); PAR, private allele richness (standard error of the mean); D, Tajima's D; H, Fay and Wu's standardised H; LD50, distance (in bp) at which linkage disequilibrium reaches half of its maximum value. Only core chromosomes BCIN01 to BCIN16 were included in calculations. Per site estimates of $\pi$ , $\rho$ and $\theta$ were computed by summing across chromosomes and dividing by number of sites covered. Tajima's D and Fay and Wu's standardised H were computed across 10kb windows, and averaged. AR and PAR were computed using a generalized rarefaction approach and a standardized sample size of two haploid genomes.

The site frequency spectra estimated in populations G1 and T were U-shaped, indicating an excess of high-frequency derived alleles (Figure 2; population G2 was excluded because of too small a sample size), consistent with ongoing episodes of positive selection, mis-assignment of ancestral alleles or gene flow (Marchi & Excoffier 2020). Estimates of Tajima’s D were positive but close to zero in clusters T and G1 (T: D=0.309; G1: D=0.167), indicating a slight deficit of low frequency variants (Table 2; Supplementary Table 2), consistent with balancing selection or population contraction. In cluster G2, the estimated Tajima’s D was D=1.470 but the estimate was likely upwardly biased by the small sample size, because Watterson’s θ is underestimated when sample size is small. The distance at which linkage disequilibrium (LD) decayed to 50% of its maximum was an order of magnitude longer in G2 (LDdecay50: 16,800bp) than in T (LDdecay50: 8600bp) and G1 (LDdecay50: 3100bp) (Supplementary Figure S3). The recombination rate was higher in G2 (ρ=0.2047/bp) than in G1 and T (G1: ρ=0.0178/bp; T: ρ=0.0108/bp).

**Figure 2.**
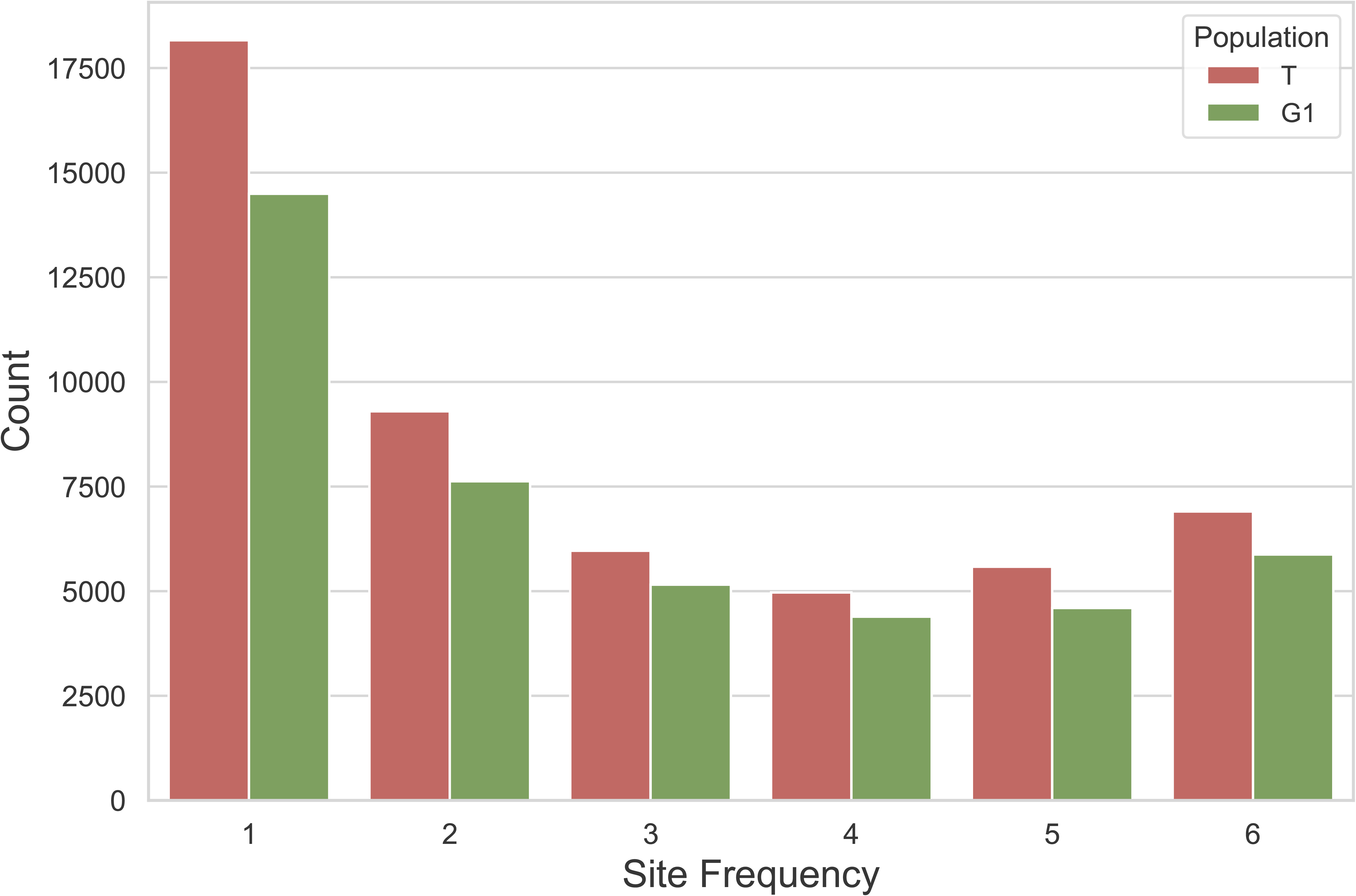
Site frequency spectra estimated in populations T2 and Vv1 based on 85,415 SNPs using the python package dadi, projecting sample sizes to seven haploid genomes. Projection consists in averaging over all possible re-samplings of the larger sample size data, thus biallelic positions with data for less than seven individuals are not included in calculations and not counted as SNPs.

The accessory chromosome BCIN18 showed no polymorphism in all three population and the accessory chromosome BCIN17 showed no polymorphism in population G2 (Supplementary Table S2). However, it should be noted that coverage analysis also revealed that these accessory chromosomes, which contain a reduced number of genes (23 and 19 respectively, van Kan *et al.* 2017), were distributed irrespectively of the host of origin: BCIN17 was found present in all but three isolates (Sl9, Sl10 and Vv15), while BCIN18 was only present in five isolates (Sl3, Sl5, Sl9, Vv9 and Vv11; Supplementary Table S3). In population T, the accessory chromosome BCIN17 displayed a relatively high and positive value of Tajima’s D (D=1.37; Supplementary Table S2) and approximately twice as much nucleotide diversity as in core chromosomes (π=0.0058/bp; Supplementary Table S2). In population G1, accessory chromosome BCIN17 displayed a negative value of Tajima’s D (D=- 0.845) and two orders of magnitude less nucleotide diversity than core chromosomes (π=5.7e-5/bp; Supplementary Table S2). The differences in Tajima’s D estimates for BCIN17 reflect the existence of two divergent haplotypes in T, but not in G1 (not shown).

### Isolates from the T population are more aggressive on tomato plants

To test whether isolates collected on tomato are more aggressive on their host of origin, compared to isolates collected on grapevine, pathogenicity assays were performed on whole tomato plants in controlled conditions. To assess differences among isolates in terms of aggressiveness on grape and tomato, we used a non-parametric analysis of variance, considering the average values for each of the independent pathogenicity tests as replications. We found a significant isolate effect (Kruskal Wallis test, H (28, N= 87) =69.89221, p < 0.0001), consistent with the wide range of aggressiveness levels observed for the 29 isolates.

To compare the aggressiveness on tomato of isolates from different hosts of origin (tomato vs. grape), we tested for differences in the distribution of the aggressiveness index between isolates originating from the two types of hosts, considering the independent repetitions of the pathogenicity test as blocks and the 13 isolates from tomato and 16 isolates from grapevine as replicates. We observed a significant effect of the host of origin (Figure 3, Mann-Whitney U test, p < 0.0001) and T isolates collected on tomato were on average 2.7 times more aggressive on tomato plants than isolates collected on grapevine.

**Figure 3.**
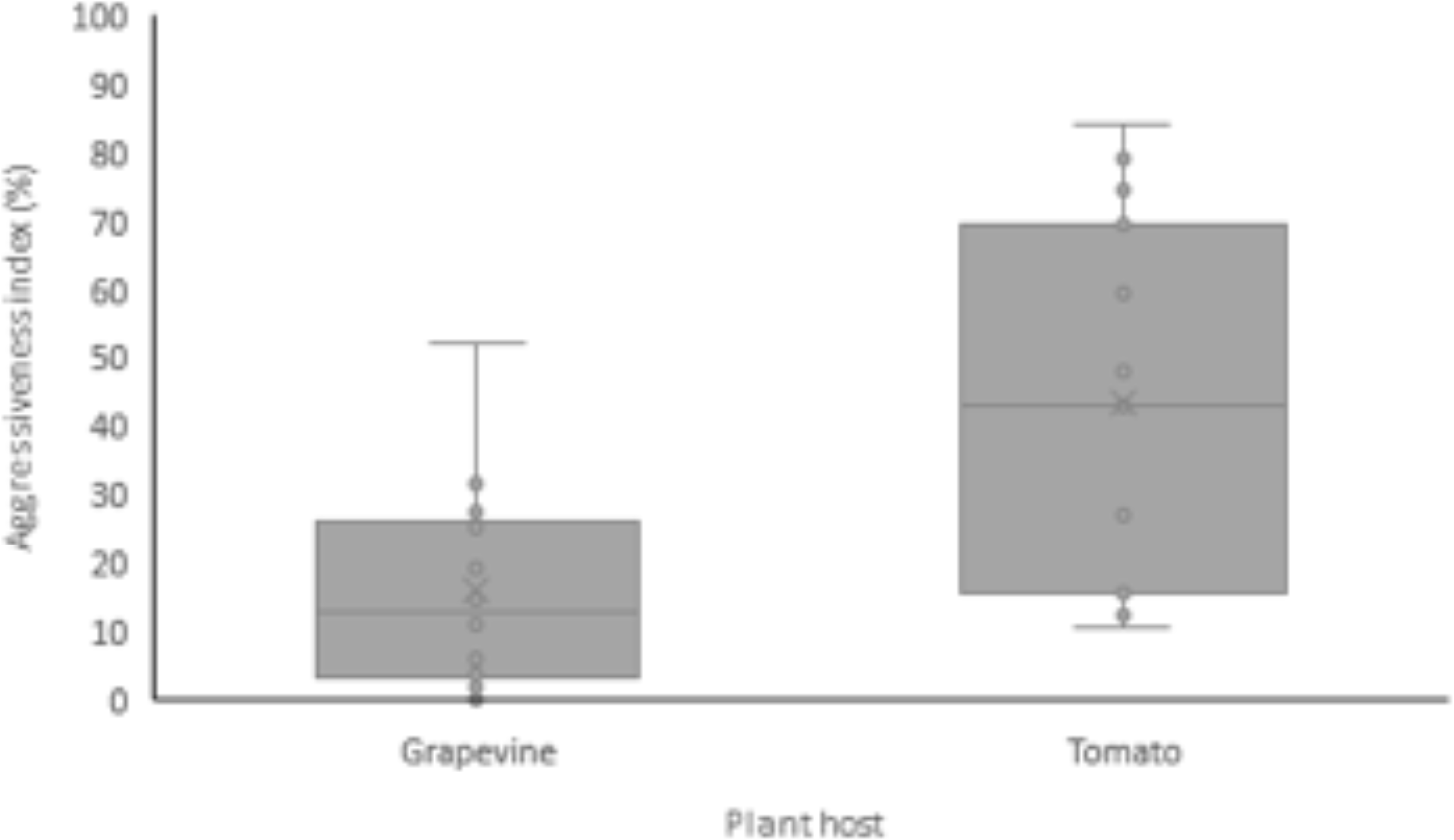
Boxplots representing the extent of aggressiveness (% relative to the reference isolate BC1) on tomato of the *Botrytis cinerea* isolates collected on grapevine (n=16) and tomato (n=13). For each boxplot, mean (crosses), median (horizontal lines), values of the aggressiveness index (circles), 25-75% quartiles, and maximum and minimum values are represented.

To compare the aggressiveness on tomato of isolates from three different clusters (T, 13 isolates; G1, 12 isolates; G2, 4 isolates), we tested whether the aggressiveness index of isolates from different clusters originate from the same distribution. A Kruskall-Wallis test rejected the null hypothesis that all clusters display the same median of the aggressiveness index (H (2, N= 87) =19.80083, p < 0.0001). The T population was significantly different from the G1 population but not from G2 (Kruskall-Wallis test, p< 0.0001 and p = 0.37, respectively). G2 and G1 were not significantly different (Kruskall-Wallis test, p = 0.57).

### Gene content slightly differs between tomato- and grapevine-associated populations

As variation in gene content can be involved in adaptation to novel hosts (Cummings *et al.* 2004; Inoue *et al.* 2017; Langridge *et al.* 2015), we sought to identify genes specific to the tomato (T) and grapevine (G1 and G2) populations. We first explored the mapping coverage of genes previously identified in the B05.10 reference to identify sets of genes that were missing or showing duplication events (Supplementary Table S4). Five B05.10 genes were identified as missing in the T population, including four consecutive genes in the subtelomeric region of chromosome BCIN02 that could correspond to a secondary metabolism gene cluster. Among these four genes, one is coding for a NRPS-like enzyme similar to the protein MelA of *Aspergillus terreus* involved in the biosynthesis of an α-keto acid dimer (Geib *et al.* 2016), two other genes encode putative biosynthetic enzymes (FAD-binding and enoyl reductase domains). Mapping coverage of B05.10 genes also suggested some possible duplication events in a subtelomeric region of chromosome BCIN08, with T isolates showing approximately three times as many reads as the G1 and G2 isolates for the four consecutive genes Bcin08g00060 to Bcin08g00090. This suggested that the corresponding region of at least 25 kb would be in three copies in the genomes of the T isolates. Among the four duplicated genes, two encode carbohydrate-active enzymes (CAZymes) known as plant cell wall degrading enzymes (PCWDEs) as they act on pectin (glycoside hydrolase GH28) and hemicellulose or pectin side chains (GH43).

We also analysed the variation in gene content using a different approach that makes no use of reference genome B05.10. We built *de novo* assemblies of the genomes of T, G1 and G2 isolates. The genome assembly size of the 28 isolates ranged from 41 Mb to 42.5 Mb (Supplementary Table S5), which was slightly smaller than the genome assembly size of the B05.10 reference isolate (42.6 Mb). We then predicted genes *ab initio* using FGenesh. The number of predicted genes ranged from 11,109 to 11,311 among genomes. To compare gene content in the T, G1 and G2 populations, we used Orthofinder to identify 12,319 groups of orthologous sequences (i.e., orthogroups). The number of groups of orthogroups shared by pairs of isolates within populations was higher that between populations (Supplementary Table S6). By looking for orthologous groups that were present in at least 75% of the genomes of a focal population and missing in other populations, we identified 21 G1-specific genes, a single G2-specific gene, five genes specific to the G1 and G2 populations (those already detected with the first approach described above), two genes missing specifically in the G1 population, and a single gene specific to the T population (Supplementary Table S7). This latter gene was a GH71 glycoside hydrolase (OG0011490, an α-1,3-glucanase; (Lombard *et al.* 2014) acting on fungal cell wall. Among the genes specific to G1, we found a GH10 glycoside hydrolase (in OG0011469; (Lombard *et al.* 2014) acting on plant cell wall (*i.e.* hemicellulose). The proteins encoded by the other G1-specific genes had no functional prediction though four of them shared a domain typical of metalloenzymes (IPR11249) with putative peptidase activities and three other ones showed a versatile protein-protein interaction motif involved in many functions (IPR011333). Two proteins with secretion signal peptides were also found specific to G1 (OG0011305 and OG0011366), with OG0011366 having a predicted function of interferon alpha-inducible protein-like (Rosebeck & Leaman 2008) and a predicted transmembrane helix.

Together, these analyses revealed that the magnitude of gene content variation is limited between *B. cinerea* populations, which emphasizes the need to investigate differences in allelic content at shared genes for elucidating the genomic basis of host specialization.

### McDonald-Kreitman tests of positive selection identify genes related to virulence

We investigated differences in the direction and intensity of natural selection driving the evolution of gene sequences in the two populations with the greatest difference in terms of quantitative pathogenicity on grape and tomato, which are also the two populations with the largest sample size (G1 and T). More specifically, we searched for genes with signatures of positive selection in both populations that also show high sequence divergence between populations, or genes with signatures of positive selection in one population, but not in the other population. The direction and intensity of selection was estimated using neutrality indexes computed for each individual gene in each population based on McDonald-Kreitman tables of polymorphisms and substitutions at synonymous and non-synonymous sites, using *B. fabae* as the outgroup. The neutrality index is expected to be below one for genes under positive selection (due to an excess of non-synonymous substitutions) and above one for genes under negative selection (due to a deficit of non-synonymous polymorphisms). To identify genes potentially involved in host specialization, we first selected genes showing low values of the neutrality index in both populations (log [neutrality index] ≤-0.5), and high values of the inter-population dN/dS ratio (dN/dS in the top 5% percentile; Supplementary Figure S4). This analysis identified five genes: Bcin02p04900, Bcin07p02650, Bcin09p06530, Bcin14p01690, Bcin09p02190 (Supplementary Table S8). Two genes (Bcin07p02650, Bcin14p01690) code for glycosyl hydrolases of the GH5 family, a family that includes enzymes acting on plant cell walls and enzymes act on fungal cell walls. One of the two genes (Bcin14p01690), has a cellulose binding domain which strongly suggests a role as PCWDE. Two genes (Bcin02p04900, Bcin09p02190) are involved in basic cell functions (a DNA nuclease and a protein involved in ribosome biogenesis). The last gene (Bcin09p06530) has no known domain.

We also selected genes showing low values of the neutrality index in one population (log [neutrality index] ≤-0.5), and high values in the other (log [neutrality index]>=0) (Supplementary Figure S4). This analysis identified 392 genes in G1 and 428 genes in T (Supplementary Table S8). Functional enrichment analyses revealed contrasting results in the G1 and T populations (Supplementary Table S8). In the G1 population, we identified a significant two-fold enrichment in transporter-encoding genes among the 392 genes with signatures of positive selection. These 28 transporters included many candidates with putative roles in nutrition such as the transport of sugars and amino-acids (five genes of each). Fifteen of them were proteins of the major facilitator superfamily (MFS). The MFS transporter-encoding genes included five putative sugar transporters and ten unknown transporters that could have roles in various processes, including obtaining nutrients from the host, efflux of fungi-toxic compounds or the export of fungal phytotoxins (Hartmann *et al.* 2018; Maruthachalam *et al.* 2011; Perlin *et al.* 2014). Finally, five of the 392 genes encoded ATPase transporters including the BcPrm1 P-type Ca2+/Mn2+-ATPase that mediates cell-wall integrity and virulence in *B. cinerea* (Plaza *et al.* 2015).

In the T population, the list of 428 genes with signatures of positive selection showed a significant 2.5-fold enrichment in genes encoding for proteins involved in oxidative stress response (eight genes). These genes encode enzymes that are able to detoxify reactive oxygen species *i.e.* glutathione-S-transferases (BcGST1, 9 and 24), the superoxide dismutase BcSOD1, and two peroxidases (BcPRX8 and BcCCP1). In addition, the list of 428 genes also included *BcatrO*, which encodes the transporter BcAtrO involved in the resistance to H_2_O_2_ (Pane *et al.* 2008).

A 2.5-fold enrichment was also observed for the genes coding for CAZYmes acting as PCWDEs (ten genes) especially for those involved in the modification of hemicellulose (five genes; Espino *et al.* 2010; Supplementary Table S8) such as the xylanase BcXyn11A that is required for full virulence on tomato (Brito *et al.* 2006).

### Selective sweeps in regions encompassing genes encoding enzymes involved in carbohydrates metabolism

To identify genomic regions with signatures of selective sweeps, we conducted three different genome scans, using different features of the data: i) hapFLK, which detects hard and soft sweeps based on patterns of differentiation between clusters of haplotypes between populations; ii) *nS_L_*, which detects hard and soft sweeps based on the distribution of fragment length between mutations and the distribution of the number of segregating sites between pairs of chromosomes; iii) and SWEED’s CLR, which detects hard sweeps based on the site frequency spectrum. The *CLR* and *nS_L_* metrics are population-specific and were computed for each population independently, while the hapFLK metric is F_ST_-based and was thus computed for populations G1 and T (Supplementary figure S5). We identified candidate SNPs located in (hard or soft) selective sweeps as the SNPs that were in the top 5% of the hapFLK statistic, but also in the top 5% of either the nSL or CLR statistic in a least one population (Figure 4). In total, this approach identified 4,667 SNPs of which 1,300 were localized in coding sequences, 256 in introns, 830 in untranslated transcribed regions, and 465 less than 1500bp upstream of coding sequences. These 2,851 SNPs corresponded to 351 genes, of which 15 were identified by SNPs in the top 5% of selective sweep metrics in both populations, 200 by SNPs in the top 5% of selective sweep metrics in population T, and 175 by SNPs in the top 5% of selective sweep metrics in population G1 (Supplementary table S9).

**Figure 4.**
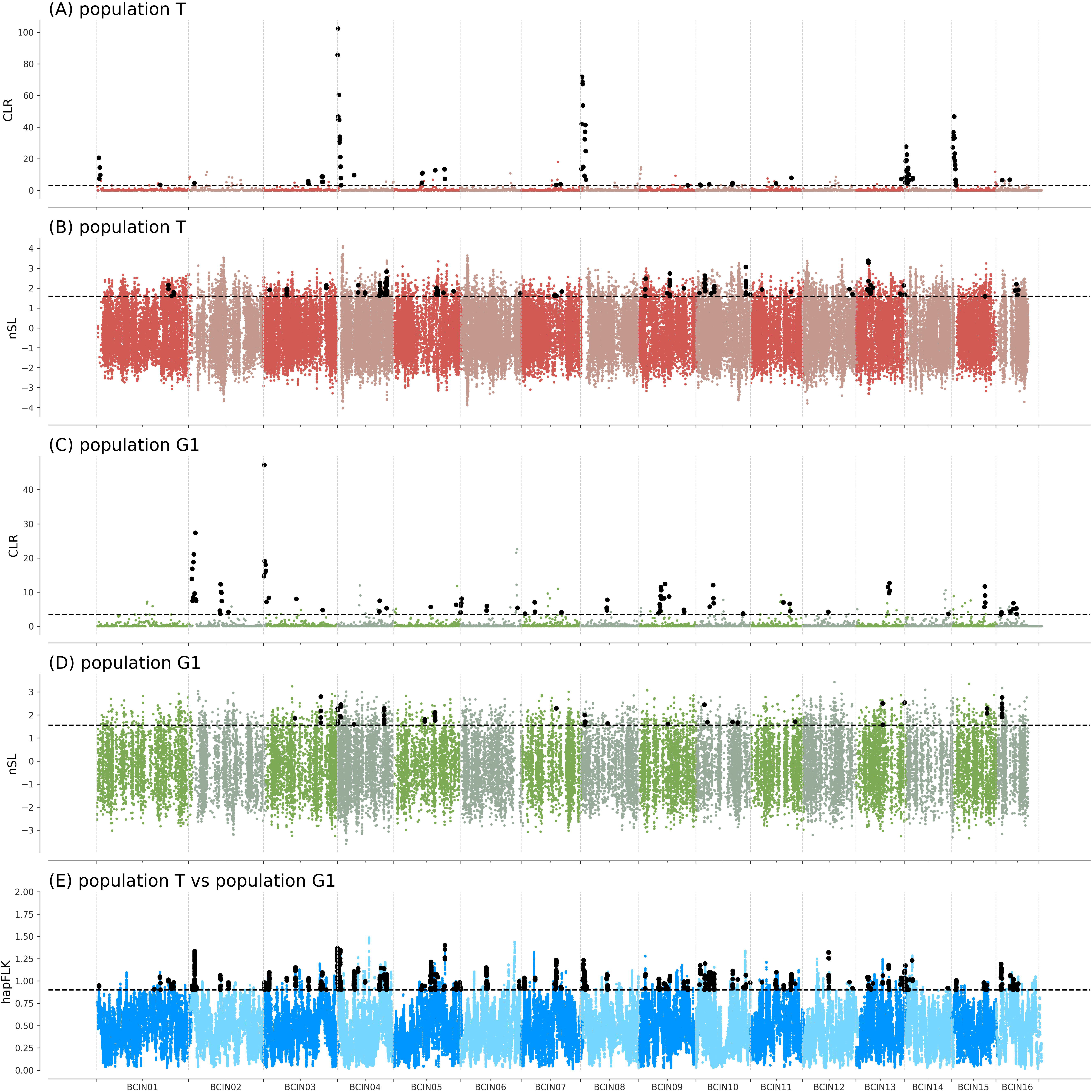
Genome scans for selective sweeps in the T and G1 populations of *Botrytis cinerea*, parasitizing tomato and grapevine, respectively. (A) and (C): Composite likelihood ratio (CLR) estimated using the SweeD software (Nielsen *et al.* 2005; Pavlidis et al., 2013) in T and G1, respectively. (B) and (D) Number of segregating sites by length, *nS_L_*, estimated using the nSL software (Ferrer-Admetlla *et al.* 2014) in T and G1, respectively. (E) hapFLK statistic estimated using the HapFLK software (Bonhomme *et al.* 2010; Fariello *et al.* 2013). Horizontal dashed black lines represent the top 5%. Vertical dashed grey lines represent the boundaries of the 16 chromosomes and the chromosome names are set along the x axis. SNPs in chromosomes with names ending with odd numbers are represented in dark colors, while SNPs in chromosomes with names ending with even numbers are represented in light colors. Black dots represent SNPs belonging to the following set: [(top 5% CLR population G1) ⋃ (top 5% CLR population T) ⋃ (top 5% nSL population G1) ⋃ (top 5% nSL population T)] ⋂ [top 5% hapFLK].

Candidate genes in the selective sweep regions of each population included genes coding for proteins with functions consistent with a role in infection, such as transporters, CAZymes, putative effectors and two genes that are confirmed virulence factors. Indeed, one region identified in the G1 population contains the gene encoding the Pectin Methyl Esterase BcPme1 required for full virulence of *B. cinerea* on several host including grapevine (Valette-Collet *et al.* 2003), and another region contains the gene encoding BcCgf1, a small secreted protein that is essential for infection structure development (Zhang *et al.* 2020). Nevertheless, it is unlikely that all genes in selective sweep regions have been direct targets of positive selection, most of them being possible hitch-hikers. This could be the reason why no significant functional enrichment was detected among these genes.

Finally, comparison of the lists of genes identified in the selective sweep regions and those with signatures of positive selection according to McDonald-Kreitman tests identified nine genes in common for the T population and four genes in common for the G1 population (Supplementary table S9). This suggests that these genes were subjected to both recurrent positive selection for amino-acid changes and to recent positive selection in populations of *B. cinerea*. In addition, we can hypothesize that these genes may be the actual targets of positive selection, and that surrounding candidate genes could be only hitch-hikers. Functional annotations of the genes that show both recurrent positive selection for amino-acid changes and recent positive selection signals further indicated a cutinase-encoding gene (Bcin01g09430) in the T population and a sugar transporter encoding gene (Bcin16g00530) in the G1 population, and also pointed out various other functions.

## Discussion

### *Genetic differentiation in* B. cinerea *between populations associated with grapevine and tomato*

Genetic structure associated with the host of origin in *B. cinerea* has been extensively investigated (reviewed in Walker 2016). Most of the studies based on sufficient sampling size (n>100) found significant population differentiation between populations of *B. cinerea* from different hosts (e.g. in Chile, Tunisia, Hungary, United Kingdom; Walker 2016). One noticeable case is the population of *B. cinerea* collected from various hosts in California (Ma & Michailides 2005), for which whole-genome sequencing data did not detect any host-associated population structure despite differences in pathogenicity against different hosts (including tomato) in cross-infectivity assays (Atwell *et al.* 2015; Caseys *et al.* 2020; Soltis *et al.* 2019). In France, previous research concluded that *B. cinerea* populations were differentiated according to some of their hosts, including tomato, grapevine and, to a lesser extent, bramble (Fournier & Giraud 2008; Walker *et al.* 2015). In two recent studies comparing the aggressiveness of isolates coming from diverse hosts, the disease severity caused by isolates from tomato was significantly greater than the severity caused by isolates from grape or other crops, thus indicating that *B. cinerea* populations parasitizing tomato were specialized to this host (Bardin *et al.* 2018; Mercier *et al.* 2019). Here we show that populations parasitizing tomato and grapevine are subdivided into three populations, two being associated with grapevine (G1 and G2) and one with tomato (T). This pattern of population genetic structure differs from previous findings (Walker et al., 2015), as three populations parasitizing grapevine had previously been detected, but studies differ in terms of sampling and genotyping schemes (thousands of SNPs vs 8 SSR markers, and an order of magnitude of difference in the size of sample sets). The clear pattern of population subdivision found in our study also stands in sharp contrast with the lack of host- or geography-associated population subdivision across various hosts, including tomato and grape, in California (Atwell *et al.* 2015; Atwell *et al.* 2018). This difference in population structure between France and California indicates that the factors leading to host-specific differentiation between *B. cinerea* pathogens from grape and tomato do not operate everywhere. The differences in LD decay (up to an order of magnitude longer in our study compared to Californian *B. cinerea*) and nucleotide diversity (half as much in our study compared to Californian *B. cinerea*) also suggest that the demographic history and population biology of the pathogen is contrasted between the two regions.

Multiple factors can contribute to reduce gene flow between populations parasitizing grapevine and tomato. A first possible factor limiting gene flow is adaptation to host. Mating in *B. cinerea* occurs on the host after infection, between individuals that were thus sufficiently adapted to infect the same host, which induces assortative mating with respect to host use and reduce opportunities for inter-population crosses (Giraud 2006; Giraud *et al.* 2010; Giraud *et al.* 2008). Another factor possibly limiting gene flow between populations infecting tomato and grape is habitat isolation (i.e. reduced encounters caused by mating in different habitats). Tomatoes are grown in nurseries before being dispatched to the fields, tunnels or greenhouses, and this may generate habitat isolation if sexual reproduction in the pathogen occurs in nurseries for the tomato-infecting population of *B. cinerea*. Such habitat isolation may contribute to promote adaptation to tomato, by preventing the immigration of alleles which are favorable for infection of non-tomato hosts but not favorable for infection of tomato. Differences in the timing of epidemics are unlikely to contribute to this habitat isolation, as the period of infection of greenhouse tomatoes runs from late to early winter, which includes the period of infection of grape. The same goes for the location of epidemics, since the sites studied were chosen because the two types of crops are grown nearby. A final possibility to explain the lack of gene flow between *B. cinerea* populations from grape and tomato is that the frequency of sexual reproduction might be lower in populations infecting greenhouse tomatoes. Higher winter temperatures and the removal of plant residues in the greenhouse represent conditions that are less conducive to sexual reproduction. However, our estimates of LD decay and recombination rates are not consistent with a substantially lower frequency of sexual reproduction in the population associated with tomato, compared to the population associated with grape.

The differences in population structure observed between *B. cinerea* populations from France and California could be due to different cultivation practices. In California, differences in pathogenicity between tomato and other hosts did not lead to genome-wide differentiation, indicating that gene flow occurs between hosts. In France, on the contrary, differences in virulence are associated to genome-wide differentiation, indicating restriction of gene flow. These differences in structure may be explained by the cultivation of tomatoes in open fields in California, which favors dispersal to other crops, while French tomatoes are generally grown in plastic tunnels or greenhouses.

### Widespread signatures of selection along genomes

We identified little variation in the gene content among T, G1 and G2 populations, with one gene specific to T, 22 genes specific to G1 and five genes shared between G1 and G2 but not T, suggesting that gene gain or loss is not the main process of adaptation to tomato. In parallel to our analysis of presence/absence variation, our genome scans for positive selection pinpointed several genomic regions which may harbour determinants of ecological differentiation between the population specialized to tomato and the population parasitizing grapevine. In order to cover multiple time scales and different signatures of positive selection, we used a variety of analytical approaches. The McDonald-Kreitman test focuses on genes and detects repeated episodes of selective sweeps fixing non-synonymous substitutions, thus generating a higher ratio of amino acid divergence to polymorphism (Dn / Pn), relative to the ratio of silent divergence to polymorphism (Ds / Ps), than expected under neutrality. The values of nucleotide diversity and Tajima’s D measured in the T population specialized in tomatoes were very close to the values measured for the two other populations, which is not consistent with a very recent origin of this population and justifies the use of the McDonald-Kreitman test. Genome scans for selective sweeps detect more recent events, and by nature these methods can also detect genes that are not directly the target of selection, but may have hitch-hiked due to physical linkage with sites under positive selection. However, the LD decay values measured for the T population remain moderate (9kb), and we used a combination of different selective sweep metrics to substantially shorten the list of candidate genes, which should reduce the impact of genetic hitch-hiking on our list of genes under recent positive selection. Despite using a more conservative approach, we identified more selective sweeps than in the generalist *Sclerotinia sclerotiorum* fungus (Derbyshire *et al.* 2019), or in the Californian generalist population of *B. cinerea* (Soltis *et al.* 2019).

### Genes under positive selection

We identified a number of genes showing signatures of positive selection using the approaches discussed above and highlighting potential candidates for their role in host specialization. Functional annotation of the *B. cinerea* genome and previous experimental studies provided lists of genes involved in host-pathogen interaction and in other developmental processes (Amselem *et al.* 2011; Nakajima & Akutsu 2014; Rodriguez-Moreno *et al.* 2018). We used these published lists of genes to investigate whether some specific biological processes were subjected to positive selection in the different populations. Our data revealed that the five genes showing the strongest signatures of selection in both the T and G1 populations, but also showing high sequence divergence between the two populations, included two genes encoding for CAZymes with glycoside hydrolase activity, which are potential PCWDEs. We also found that the 428 genes showing the strongest signatures of selection in the T population with McDonald-Kreitman tests were enriched both in genes coding for PCWDEs and in genes coding for enzymes involved in the oxidative stress response. Notably, ten genes encoding secreted CAZymes targeting compounds of the plant cell wall, *i.e.* cellulose and pectin (Amselem et al., 2011; Lombard *et al.* 2014), were found under positive selection. One of these ten genes encodes the xylanase BcXyn11A that has previously been shown to be a virulence factor on tomato (Brito *et al.* 2006). Another one encodes a cutinase (Bcin01g09430) that was further detected in a selective sweep region suggesting, recurrent and recent positive selection events. Additional PCWDEs were found in other genomic regions identified as selective sweeps. Finally, our comparative genomic analysis suggested that a subtelomeric region that contains two PCWDEs acting on pectin and/or hemicellulose is duplicated in the T population. Necrotrophic species have important repertoires of CAZymes especially those corresponding to PCWDEs which are known to act as major virulence factors in fungi (Zhao *et al.* 2013; Rodriguez-Moreno *et al.* 2018). The genome of the reference isolate of *B. cinerea* (B05.10) revealed 118 PCWDEs (Amselem et al., 2011) and our data suggest that, within this repertoire, some cellulases and pectinases may be of particular importance for the degradation of tomato cell wall. SNPs within a pectinesterase gene were also associated with virulence on tomato in a previous genome-wide association study (Soltis *et al.* 2019).

In addition to PCWDEs, the single gene that was present in the T population but missing in the G1 and G2 populations encoded a CAZyme acting on the fungal cell wall, an α-1,3-glucanase classified as a member of the GH71 family. In the fungal cell wall, α-1,3-glucan is a major component that encloses the α-(1,3)-glucan-chitin fibrillar core. Because of its external localization and specific composition, α-1,3-glucan of pathogenic fungi plays a major role in infection-related morphology and host recognition (Beauvais *et al.* 2013; King *et al.* 2017). A dozen of genes of *B. cinerea* encode for enzymes of the GH71 family (Amselem et al., 2011). The T-specific GH71 CAZyme might therefore have been retained in the T population as a mean to specifically facilitate infection of tomato by modification of the fungal cell wall resulting in adaptive morphological changes or impairment of host recognition.

As mentioned above, the McDonald-Kreitman tests also indicated that eight genes coding for enzymes that detoxify reactive oxygen species showed signatures of positive selection in the T population. During infection, *B. cinerea* encounters an oxidative burst, an early host response that results in the death of plant cells. This mechanism is used in turn by *B. cinerea* to achieve full virulence but this also implies that the fungus has to resist to this toxic environment. The fungal oxidative stress response system includes detoxifying enzymes such as superoxide dismutases (SODs) that convert O_2_ into the less toxic H_2_O_2_, as well as catalases, peroxidases and peroxiredoxins that convert H_2_O_2_. Additional non-enzymatic mechanisms include the oxidation of compounds such as glutathione (Heller & Tudzynski 2011). In addition to the genes encoding SOD (BcSOD3), peroxidases (BcCCP1), peroxiredoxins (BcPRX8) and other detoxifying enzymes sur as glutathione-S-transferases (BcGST1, 9 and 24), a gene encoding the transporter BcAtrO also showed a signature of positive selection in the T population. Inactivation of this gene previously suggested that it allows the efflux of H_2_O_2_ and resistance this reactive oxygen species (Pane *et al.* 2008). Altogether, our data suggest that the oxidative burst occurring in the *B. cinerea*/tomato interaction is particularly challenging for the fungus.

### Concluding remarks

We identified a population of *B. cinerea* specialized to tomato, which diverged from a grapevine-associated population. Genome scans for selective sweeps and McDonald-Kreitman tests revealed widespread signatures of positive selection that identified genes that may contribute to the pathogen’s adaptation to its tomato host. Candidate genes for specialization to tomato were significantly enriched in those encoding cellulases, pectinases and enzymes involved in the oxidative stress response, suggesting that the ability to degrade the host cell wall and to cope with the oxidative burst are two key process in the *B. cinerea*/tomato interaction. Our work sets the stage for future studies aiming to elucidate the phenotypic and fitness effects of the candidate genes for specialization of *B. cinerea* to tomato, for instance by knocking-out or replacing candidate genes for host specialization.

## Acknowledgements

AM was supported by a grant from the Doctoral School « Sciences du Végétal », Université Paris-Saclay. We are grateful to INRAE-LIPM Bioinformatics Platform (Jérôme Gouzy, Sébastien Carrère, Erika Sallet, Ludovic Legrand) and Genotoul Bioinformatics Platforms Toulouse Occitanie (Bioinfo Genotoul, doi: 10.15454/1.5572369328961167E12) for providing computational support and resources. This work was supported by a grant overseen by the French National Research Agency (ANR) as part of the “Investissements d’Avenir” Programme (LabEx BASC; ANR-11-LABX-0034) and by the INRAE department SPE. The BIOGER unit also benefits from the support of “Saclay Plant Science-SPS” (ANR-17-EUR-0007).

