## Supplementary table S3 for "Population Genomics Reveals Molecular Determinants of Specialization to Tomato in the Polyphagous Fungal Pathogen *Botrytis cinerea* in France"

**Supplementary Table S3.** Coverage of *Botrytis cinerea* B05.10 mini-chromosomes BCIN17 (=Chr17) and BCIN18 (=Chr18) by the Illumina reads of the libraries from the G1, G2 and T populations.

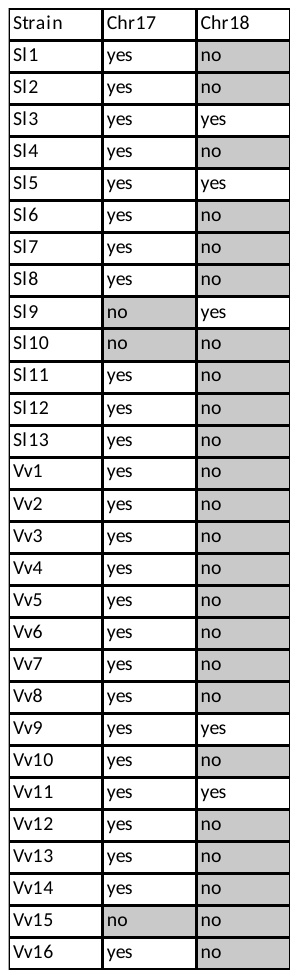
