## Supplementary table S6 for "Population Genomics Reveals Molecular Determinants of Specialization to Tomato in the Polyphagous Fungal Pathogen *Botrytis cinerea* in France"

**Supplementary Table S6.** Number of orthogroups shared between each pair of individuals of the T, G1 and G2 populations. Cells are colored according to the number of shared orthogroups, with darker shades closer to the maximum number of orthogroups in a unique individual (11219; Vv14) and lighter shades closer to the minimum value of shared orthogroups between two individuals (10612; between Sl6 and Vv6).

|  | Vv1 | Vv2 | Vv3 | Vv4 | Vv5 | Vv6 | Vv7 | Vv10 | Vv12 | Vv13 | Vv14 | Vv16 | Vv8 | Vv9 | Vv11 | Vv15 | Sl1 | Sl2 | Sl3 | Sl4 | Sl5 | Sl6 | Sl7 | Sl8 | Sl9 | Sl10 | Sl11 | Sl12 |
| --- | --- | --- | --- | --- | --- | --- | --- | --- | --- | --- | --- | --- | --- | --- | --- | --- | --- | --- | --- | --- | --- | --- | --- | --- | --- | --- | --- | --- |
| Vv1 | 11200 | 10864 | 10833 | 10891 | 10880 | 10857 | 10830 | 10845 | 10873 | 10839 | 10891 | 10861 | 10727 | 10721 | 10720 | 10738 | 10681 | 10706 | 10704 | 10707 | 10681 | 10665 | 10676 | 10696 | 10700 | 10684 | 10716 | 10683 |
| Vv2 | 10864 | 11112 | 10802 | 10816 | 10835 | 10813 | 10788 | 10798 | 10810 | 10793 | 10808 | 10842 | 10692 | 10686 | 10700 | 10689 | 10660 | 10686 | 10681 | 10682 | 10660 | 10644 | 10649 | 10656 | 10677 | 10651 | 10683 | 10659 |
| Vv3 | 10833 | 10802 | 11140 | 10853 | 10781 | 10867 | 10763 | 10792 | 10798 | 10764 | 10853 | 10806 | 10687 | 10669 | 10670 | 10696 | 10642 | 10661 | 10679 | 10663 | 10643 | 10619 | 10641 | 10643 | 10670 | 10633 | 10675 | 10653 |
| Vv4 | 10891 | 10816 | 10853 | 11161 | 10846 | 10820 | 10795 | 10828 | 10812 | 10818 | 10881 | 10832 | 10689 | 10690 | 10688 | 10697 | 10674 | 10679 | 10670 | 10681 | 10657 | 10644 | 10647 | 10666 | 10697 | 10661 | 10694 | 10676 |
| Vv5 | 10880 | 10835 | 10781 | 10846 | 11175 | 10822 | 10801 | 10818 | 10866 | 10801 | 10831 | 10829 | 10692 | 10686 | 10687 | 10691 | 10676 | 10678 | 10684 | 10683 | 10661 | 10651 | 10654 | 10683 | 10667 | 10638 | 10677 | 10655 |
| Vv6 | 10857 | 10813 | 10867 | 10820 | 10822 | 11155 | 10769 | 10774 | 10803 | 10768 | 10843 | 10792 | 10671 | 10666 | 10686 | 10692 | 10654 | 10666 | 10660 | 10657 | 10638 | 10612 | 10637 | 10685 | 10660 | 10639 | 10680 | 10654 |
| Vv7 | 10830 | 10788 | 10763 | 10795 | 10801 | 10769 | 11128 | 10845 | 10794 | 10808 | 10813 | 10844 | 10669 | 10673 | 10671 | 10665 | 10640 | 10659 | 10655 | 10660 | 10642 | 10625 | 10623 | 10664 | 10651 | 10634 | 10670 | 10658 |
| Vv10 | 10845 | 10798 | 10792 | 10828 | 10818 | 10774 | 10845 | 11129 | 10786 | 10802 | 10849 | 10807 | 10640 | 10674 | 10664 | 10659 | 10639 | 10639 | 10656 | 10651 | 10637 | 10619 | 10629 | 10656 | 10645 | 10621 | 10671 | 10642 |
| Vv12 | 10873 | 10810 | 10798 | 10812 | 10866 | 10803 | 10794 | 10786 | 11183 | 10796 | 10841 | 10821 | 10726 | 10683 | 10701 | 10724 | 10657 | 10678 | 10682 | 10690 | 10656 | 10642 | 10654 | 10697 | 10668 | 10634 | 10686 | 10673 |
| Vv13 | 10839 | 10793 | 10764 | 10818 | 10801 | 10768 | 10808 | 10802 | 10796 | 11130 | 10833 | 10808 | 10679 | 10672 | 10682 | 10663 | 10653 | 10669 | 10657 | 10661 | 10651 | 10642 | 10634 | 10666 | 10650 | 10639 | 10690 | 10668 |
| Vv14 | 10891 | 10808 | 10853 | 10881 | 10831 | 10843 | 10813 | 10849 | 10841 | 10833 | 11219 | 10841 | 10722 | 10683 | 10691 | 10705 | 10642 | 10667 | 10655 | 10659 | 10648 | 10629 | 10637 | 10672 | 10658 | 10637 | 10685 | 10662 |
| Vv16 | 10861 | 10842 | 10806 | 10832 | 10829 | 10792 | 10844 | 10807 | 10821 | 10808 | 10841 | 11165 | 10713 | 10690 | 10712 | 10695 | 10679 | 10686 | 10690 | 10692 | 10678 | 10657 | 10650 | 10675 | 10674 | 10656 | 10700 | 10684 |
| Vv8 | 10727 | 10692 | 10687 | 10689 | 10692 | 10671 | 10669 | 10640 | 10726 | 10679 | 10722 | 10713 | 11095 | 10793 | 10779 | 10760 | 10715 | 10732 | 10737 | 10724 | 10721 | 10698 | 10706 | 10725 | 10713 | 10676 | 10738 | 10736 |
| Vv9 | 10721 | 10686 | 10669 | 10690 | 10686 | 10666 | 10673 | 10674 | 10683 | 10672 | 10683 | 10690 | 10793 | 11094 | 10828 | 10792 | 10734 | 10744 | 10780 | 10755 | 10760 | 10716 | 10735 | 10735 | 10760 | 10707 | 10773 | 10746 |
| Vv11 | 10720 | 10700 | 10670 | 10688 | 10687 | 10686 | 10671 | 10664 | 10701 | 10682 | 10691 | 10712 | 10779 | 10828 | 11113 | 10802 | 10742 | 10751 | 10780 | 10748 | 10768 | 10732 | 10740 | 10744 | 10773 | 10701 | 10785 | 10760 |
| Vv15 | 10738 | 10689 | 10696 | 10697 | 10691 | 10692 | 10665 | 10659 | 10724 | 10663 | 10705 | 10695 | 10760 | 10792 | 10802 | 11123 | 10714 | 10757 | 10746 | 10736 | 10733 | 10697 | 10704 | 10727 | 10742 | 10680 | 10754 | 10745 |
| Sl1 | 10681 | 10660 | 10642 | 10674 | 10676 | 10654 | 10640 | 10639 | 10657 | 10653 | 10642 | 10679 | 10715 | 10734 | 10742 | 10714 | 11089 | 10839 | 10844 | 10848 | 10863 | 10824 | 10797 | 10803 | 10824 | 10740 | 10803 | 10831 |
| Sl2 | 10706 | 10686 | 10661 | 10679 | 10678 | 10666 | 10659 | 10639 | 10678 | 10669 | 10667 | 10686 | 10732 | 10744 | 10751 | 10757 | 10839 | 11097 | 10853 | 10946 | 10841 | 10784 | 10842 | 10832 | 10814 | 10748 | 10807 | 10843 |
| Sl3 | 10704 | 10681 | 10679 | 10670 | 10684 | 10660 | 10655 | 10656 | 10682 | 10657 | 10655 | 10690 | 10737 | 10780 | 10780 | 10746 | 10844 | 10853 | 11132 | 10852 | 10843 | 10838 | 10851 | 10805 | 10849 | 10765 | 10844 | 10839 |
| Sl4 | 10707 | 10682 | 10663 | 10681 | 10683 | 10657 | 10660 | 10651 | 10690 | 10661 | 10659 | 10692 | 10724 | 10755 | 10748 | 10736 | 10848 | 10946 | 10852 | 11110 | 10854 | 10775 | 10828 | 10814 | 10812 | 10751 | 10806 | 10845 |
| Sl5 | 10681 | 10660 | 10643 | 10657 | 10661 | 10638 | 10642 | 10637 | 10656 | 10651 | 10648 | 10678 | 10721 | 10760 | 10768 | 10733 | 10863 | 10841 | 10843 | 10854 | 11108 | 10831 | 10798 | 10792 | 10863 | 10766 | 10819 | 10837 |
| Sl6 | 10665 | 10644 | 10619 | 10644 | 10651 | 10612 | 10625 | 10619 | 10642 | 10642 | 10629 | 10657 | 10698 | 10716 | 10732 | 10697 | 10824 | 10784 | 10838 | 10775 | 10831 | 11087 | 10769 | 10761 | 10859 | 10777 | 10804 | 10791 |
| Sl7 | 10676 | 10649 | 10641 | 10647 | 10654 | 10637 | 10623 | 10629 | 10654 | 10634 | 10637 | 10650 | 10706 | 10735 | 10740 | 10704 | 10797 | 10842 | 10851 | 10828 | 10798 | 10769 | 11074 | 10814 | 10796 | 10745 | 10806 | 10797 |
| Sl8 | 10696 | 10656 | 10643 | 10666 | 10683 | 10685 | 10664 | 10656 | 10697 | 10666 | 10672 | 10675 | 10725 | 10735 | 10744 | 10727 | 10803 | 10832 | 10805 | 10814 | 10792 | 10761 | 10814 | 11115 | 10788 | 10735 | 10807 | 10806 |
| Sl9 | 10700 | 10677 | 10670 | 10697 | 10667 | 10660 | 10651 | 10645 | 10668 | 10650 | 10658 | 10674 | 10713 | 10760 | 10773 | 10742 | 10824 | 10814 | 10849 | 10812 | 10863 | 10859 | 10796 | 10788 | 11133 | 10827 | 10808 | 10802 |
| Sl10 | 10684 | 10651 | 10633 | 10661 | 10638 | 10639 | 10634 | 10621 | 10634 | 10639 | 10637 | 10656 | 10676 | 10707 | 10701 | 10680 | 10740 | 10748 | 10765 | 10751 | 10766 | 10777 | 10745 | 10735 | 10827 | 11100 | 10756 | 10760 |
| Sl11 | 10716 | 10683 | 10675 | 10694 | 10677 | 10680 | 10670 | 10671 | 10686 | 10690 | 10685 | 10700 | 10738 | 10773 | 10785 | 10754 | 10803 | 10807 | 10844 | 10806 | 10819 | 10804 | 10806 | 10807 | 10808 | 10756 | 11128 | 10918 |
| Sl12 | 10683 | 10659 | 10653 | 10676 | 10655 | 10654 | 10658 | 10642 | 10673 | 10668 | 10662 | 10684 | 10736 | 10746 | 10760 | 10745 | 10831 | 10843 | 10839 | 10845 | 10837 | 10791 | 10797 | 10806 | 10802 | 10760 | 10918 | 11107 |
