## Supplementary table S7 for "Population Genomics Reveals Molecular Determinants of Specialization to Tomato in the Polyphagous Fungal Pathogen *Botrytis cinerea* in France"

**Supplementary table S7.** Orthogroups specific to *B. cinerea* G1, G2 or T populations.

| **Orthogroup** | **Frequency in G1 isolates** | **Frequency in G2 isolates** | **Frequency in T isolates** | **IPR annotation** | **SignalP /TMHs** | **BcinB0510** |
| --- | --- | --- | --- | --- | --- | --- |
| OG0011305 | 11 | 0 | 0 |  | yes / 0 |  |
| OG0011306 | 11 | 0 | 0 |  |  |  |
| OG0011311 | 11 | 0 | 0 |  |  |  |
| OG0011367 | 10 | 0 | 0 | IPR011333:SKP1/BTB/POZ domain superfamily; |  |  |
| OG0011376 | 10 | 0 | 0 | IPR000210:BTB/POZ domain; IPR011333:SKP1/BTB/POZ domain superfamily; |  |  |
| OG0011377 | 10 | 0 | 0 | IPR000210:BTB/POZ domain; IPR011333:SKP1/BTB/POZ domain superfamily; |  |  |
| OG0011274 | 11 | 0 | 0 |  |  |  |
| OG0011300 | 11 | 0 | 0 | IPR001214:SET domain; |  |  |
| OG0011308 | 10 | 0 | 0 | IPR008999:Actin-crosslinking; |  |  |
| OG0011399 | 10 | 0 | 0 | IPR011249:Metalloenzyme, LuxS/M16 peptidase-like; IPR032632:Peptidase M16, middle/third domain; |  |  |
| OG0011309 | 11 | 0 | 0 | IPR011249:Metalloenzyme, LuxS/M16 peptidase-like; |  |  |
| OG0011366 | 10 | 0 | 0 | IPR009311:Interferon alpha-inducible protein IFI6/IFI27-like; | yes / 3 |  |
| OG0011368 | 10 | 0 | 0 | IPR002893:Zinc finger, MYND-type; IPR038727:NadR/Ttd14, AAA domain; |  |  |
| OG0011469 | 9 | 0 | 0 | IPR001000:Glycoside hydrolase family 10 domain; IPR017853:Glycoside hydrolase superfamily; |  |  |
| OG0011378 | 10 | 0 | 0 | IPR011249:Metalloenzyme, LuxS/M16 peptidase-like; |  |  |
| OG0010863 | 11 | 0 | 0 | IPR002562:3'-5' exonuclease domain; IPR012337:Ribonuclease H-like superfamily; IPR036397:Ribonuclease H superfamily; |  |  |
| OG0011436 | 9 | 0 | 0 | IPR011249:Metalloenzyme, LuxS/M16 peptidase-like; IPR032632:Peptidase M16, middle/third domain; |  |  |
| OG0011228 | 12 | 0 | 0 |  |  |  |
| OG0011051 | 12 | 0 | 0 |  |  |  |
| OG0011046 | 12 | 0 | 0 | IPR027417:P-loop containing nucleoside triphosphate hydrolase; |  |  |
| OG0011235 | 11 | 0 | 0 | IPR022099:Protein of unknown function DUF3638; |  |  |
| OG0011110 | 11 | 0 | 0 |  |  |  |
| OG0012017 | 0 | 3 | 0 |  |  |  |
| OG0011490 | 0 | 0 | 9 | IPR005197:Glycoside hydrolase family 71; |  |  |
| OG0010991 | 12 | 4 | 0 | IPR002938:FAD-binding domain; IPR036188:FAD/NAD(P)-binding domain superfamily; |  | Bcin02g00013.1 |
| OG0010990 | 12 | 4 | 0 | IPR011032:GroES-like superfamily; IPR020843:Polyketide synthase, enoylreductase domain; IPR036291:NAD(P)-binding domain superfamily; |  | Bcin02g00014.1 |
| OG0010989 | 12 | 4 | 0 |  |  | Bcin02g00015.1 |
| OG0010997 | 12 | 4 | 0 | IPR036736:ACP-like superfamily;IPR029058:Alpha/Beta hydrolase fold;IPR020845:AMP-binding, conserved site;IPR000873:AMP-dependent synthetase/ligase;IPR042099:AMP-dependent synthetase-like superfamily;IPR009081:Phosphopantetheine binding ACP domain;IPR020806:Polyketide synthase, phosphopantetheine-binding domain;IPR020802:Polyketide synthase, thioesterase domain;IPR001031:Thioesterase; |  | Bcin02g00016.1 |
| OG0011108 | 10 | 4 | 0 |  |  | Bcin15g00010.1 |
| OG0011124 | 0 | 4 | 10 | IPR013087:Zinc finger C2H2-type; IPR036390:Winged helix DNA-binding domain superfamily; IPR042065:E3 ubiquitin-protein ligase ELL-like; |  |  |
| OG0011125 | 0 | 4 | 10 |  |  |  |
