## Supplementary figure S1 for "Population Genomics Reveals Molecular Determinants of Specialization to Tomato in the Polyphagous Fungal Pathogen *Botrytis cinerea* in France"

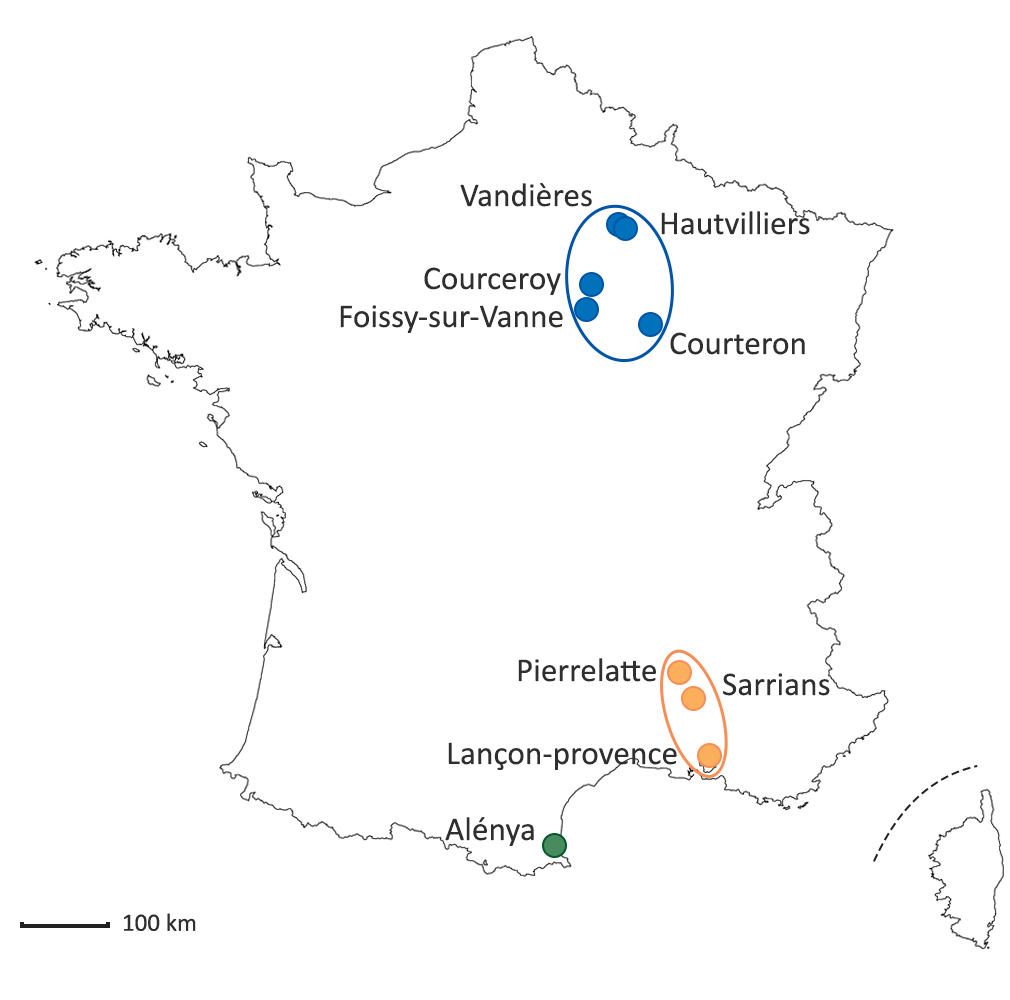

**Supplementary Figure S1.** Collection sites of *B. cinerea* pathogens sampled in France on various host plants between 2005 and 2009. Regions are represented with different colors, with Champagne in blue, Occitanie in green and Provence in orange.
