## Supplementary figure S2 for "Population Genomics Reveals Molecular Determinants of Specialization to Tomato in the Polyphagous Fungal Pathogen *Botrytis cinerea* in France"

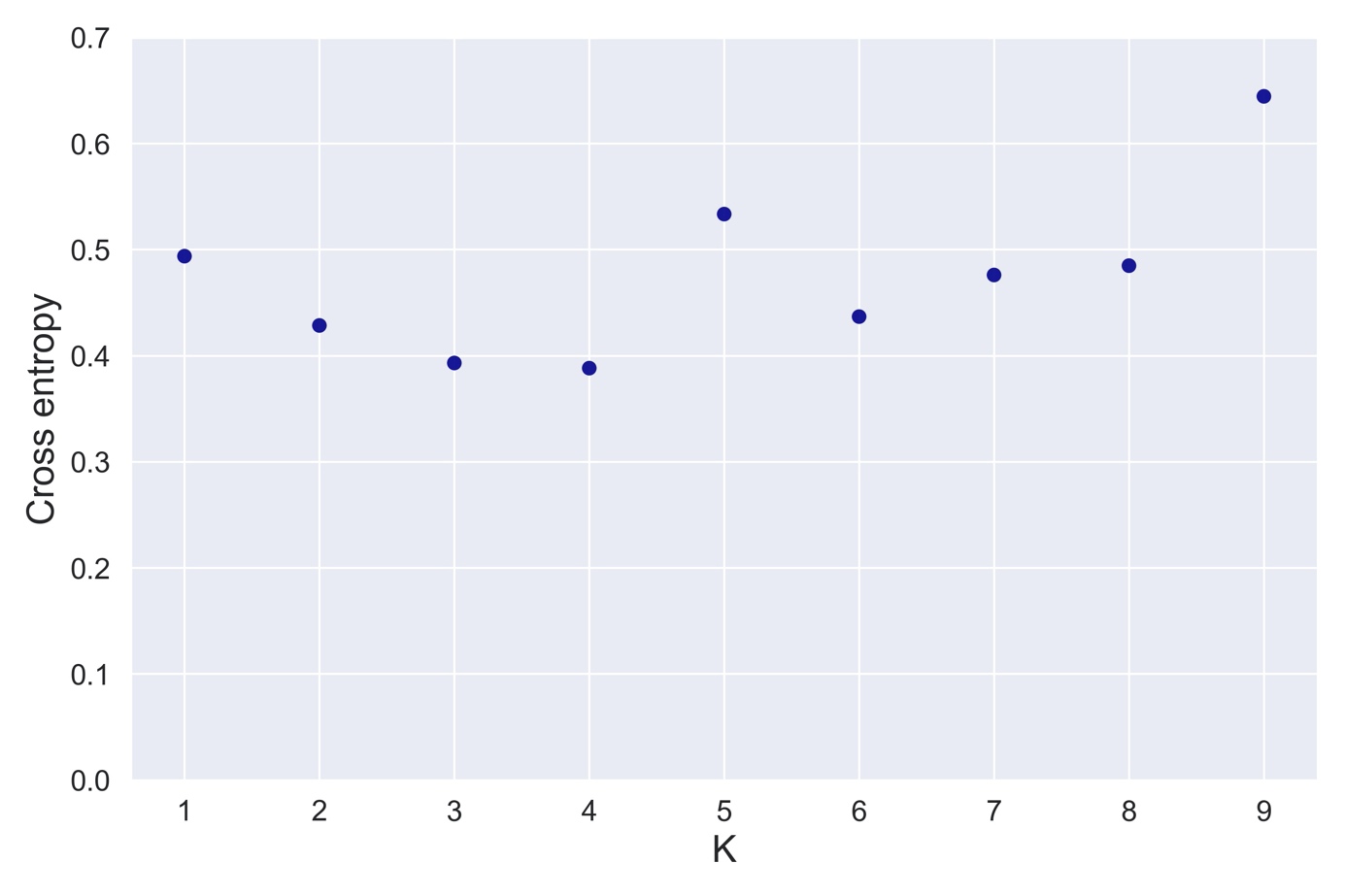

Supplementary figure S2: Cross-entropy (CE) as a function of the number of clusters K modeled in sNMF analyses of population subdivision.
