## Supplementary figure S3 for "Population Genomics Reveals Molecular Determinants of Specialization to Tomato in the Polyphagous Fungal Pathogen *Botrytis cinerea* in France"

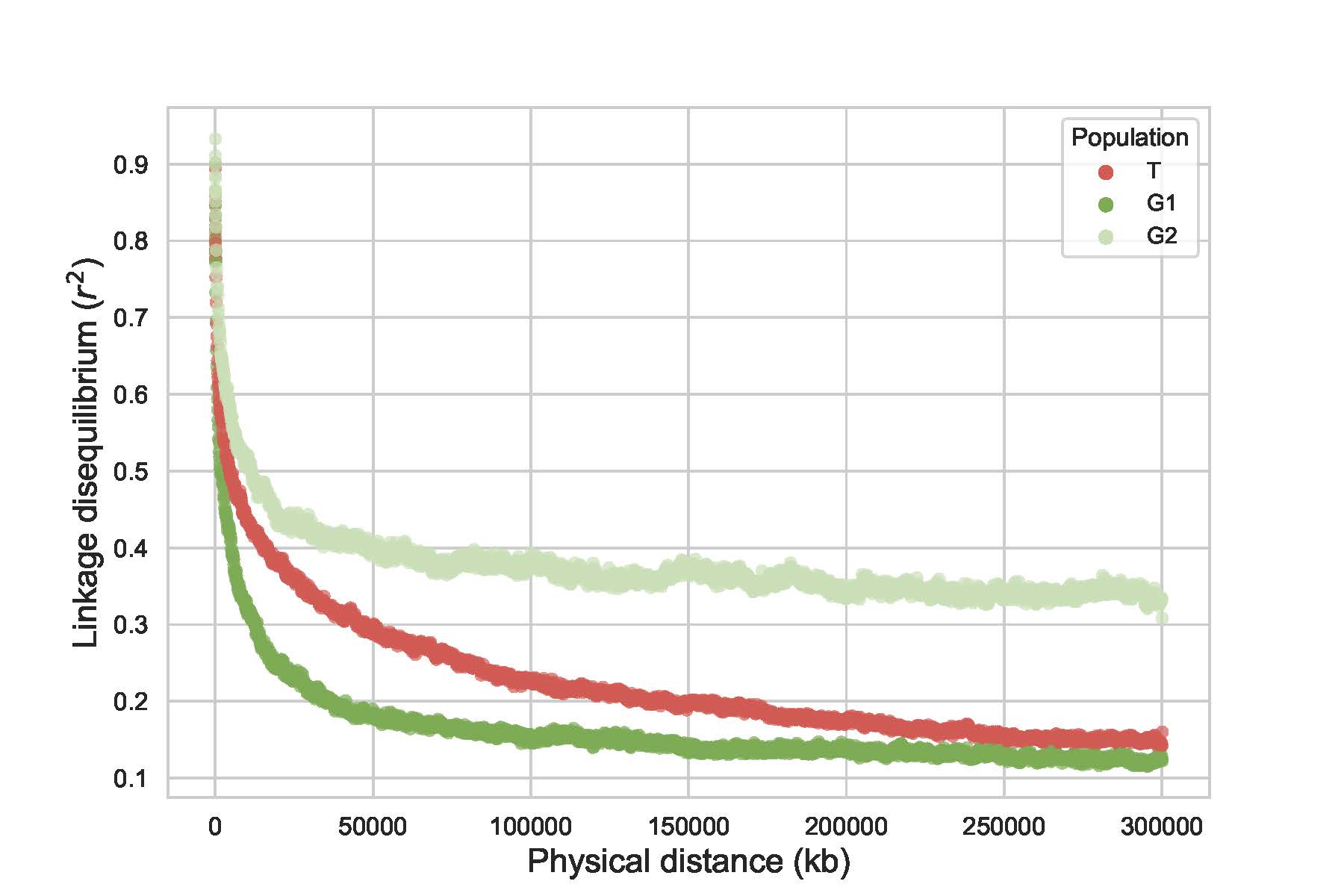

Supplementary figure S3. Linkage disequilibrium (measured as *r^2^*) as a function of the physical distance between pairs of SNPs.
