## Supplementary figure S4 for "Population Genomics Reveals Molecular Determinants of Specialization to Tomato in the Polyphagous Fungal Pathogen *Botrytis cinerea* in France"

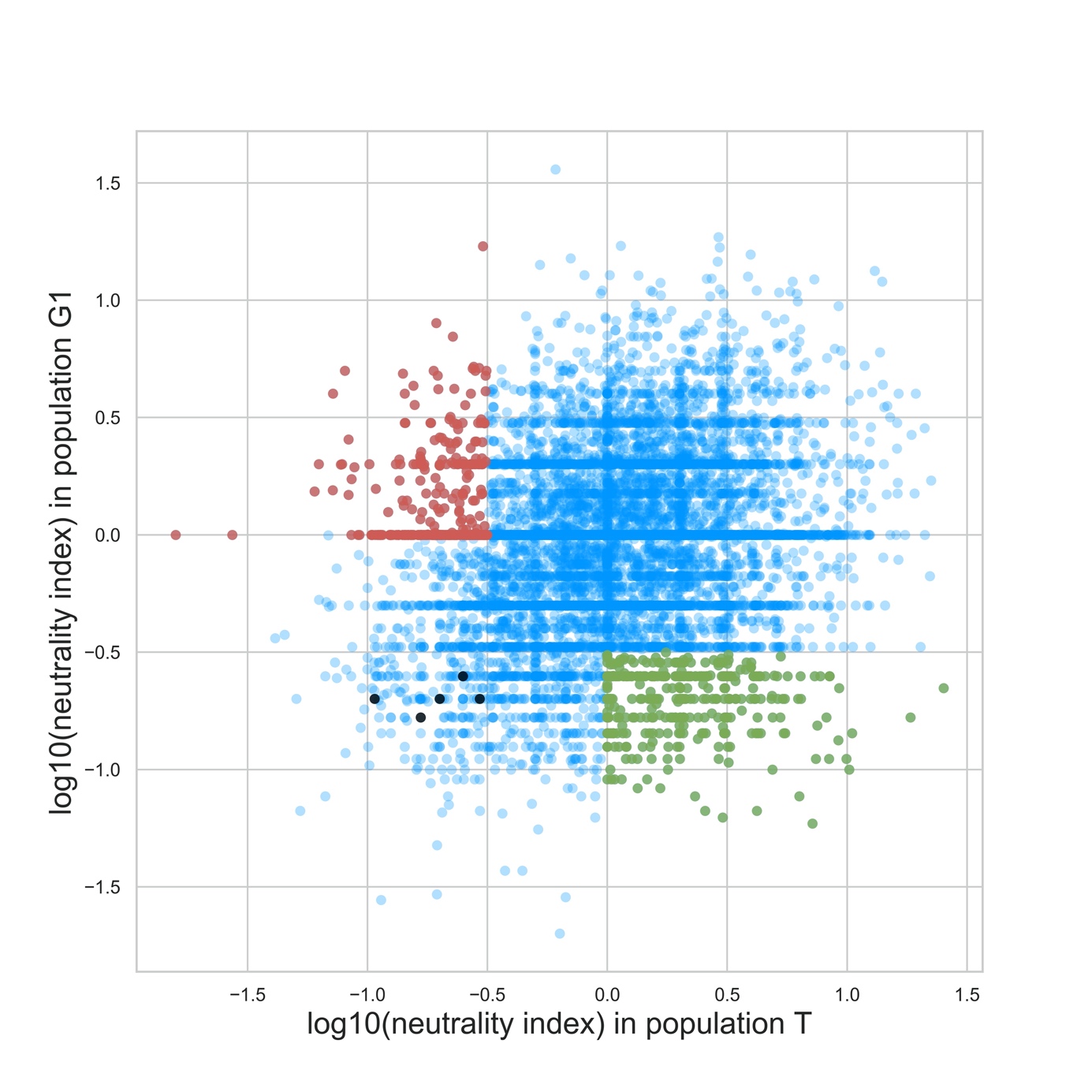

**Supplementary Figure S4.** Log_10_ of the neutrality index estimated based on polymorphism and divergence at 11,917 genes in, respectively, nine and eight isolates of the T and G1 populations of *Botrytis cinerea*. A positive Log_10_ neutrality index indicates negative selection, and a negative Log_10_ neutrality index indicates positive selection. Genes potentially involved in host specialization of the T population to tomato (red dots, top left) and of the G1 population to grape (green dots, bottom right) were identified as genes having both low neutrality index values in the focal population (log[neutrality index]<=-0.5), and high values in the alternate population (log[neutrality index]>=0).  Genes potentially involved in both host specialization of the T population to tomato and of the G1 population to grape (black dots, bottom left) were identified as genes having low neutrality index values in both populations and high dN/dS between populations (interpopulation dN/dS in the top 5% percentile).
