## Supplementary figure S5 for "Population Genomics Reveals Molecular Determinants of Specialization to Tomato in the Polyphagous Fungal Pathogen *Botrytis cinerea* in France"

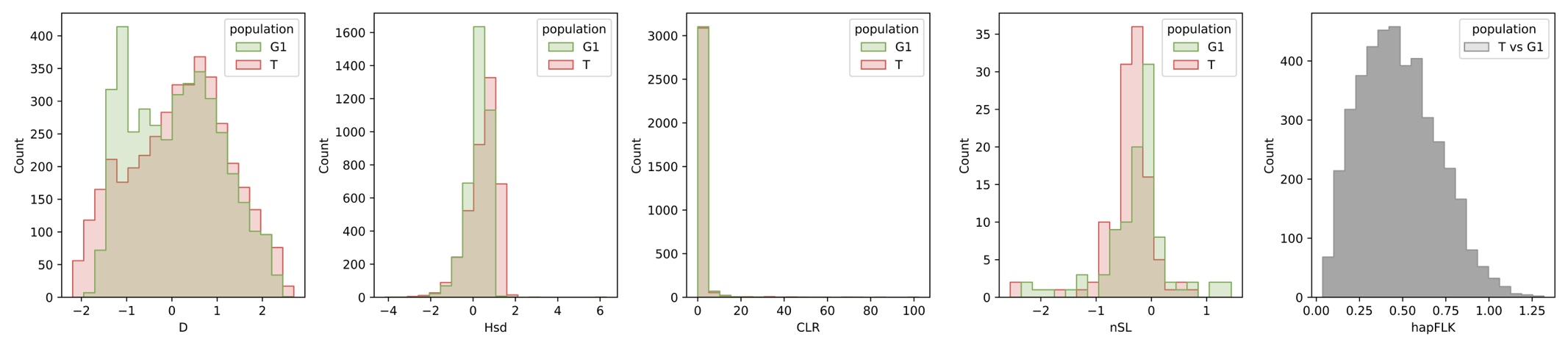

**Supplementary Figure S5.** Distribution of Tajima’s D and Fay and Wu’s standardized H (Hsd) estimated with EGGLIB 3.0, composite likelihood ratio (CLR) estimated with SWEED, nSL statistic estimated with nSL, and hapFLK statistic estimated with HAPFLK. D, Hsd, CLR and nSL were computed for each population T and G1, while hapFLK is Fst-based and was computed by contrasting population T and G1. D and Hsd were computed in 10kb genomic windows. HapFLK and nSL are computed for each SNP and were averaged in 10kb windows. CLR was computed in grids of 200 positions per chromosome.
